# Mechanisms of G-gene heterogeneities and a new method of genotyping of human metapneumovirus

**DOI:** 10.1101/2025.03.10.642527

**Authors:** Asit Kumar Chakraborty

**Affiliations:** Department of Biochemistry and Biotechnology, Vidyasagar University, Midnapore-721102, India.

**Keywords:** Metapneumovirus, genotyping, G-gene heterogeneity, termination codon mutation, RNA virus, China outbreaks, whole genome sequencing, multi-alignment and phylogram

## Abstract

Human metapneumovirus (HMPV) is similar to AMPV and HRSV single-stranded RNA viruses. Controversy of HMPV G-gene heterogeneities due to duplication, deletion, recombination and termination codon mutation (TCM) complicated genotyping of a HMPV virus since 2015. Analysis of G-proteins from complete genomes suggested ∼88% sequence homology between A and B genotypes and still sub-genotyping possible. From extensive multi-alignment studies, we concluded that the G-gene (219aa) of HMPV A-genotype (MZ504961, LC671558) was duplicated 180nt first producing G-gene with 279aa protein of A1-genotype (OL794403, OP904094,) and then 69nt deletion occurred generating G-protein with 256aa protein of A2-genotype (OP904007, MZ851793). The few amino acids lower or higher A-genotype G-proteins like 265aa (LC769214), 261aa (PV052163), 254aa (MN745086), 236aa (KC403976), 228 (KJ627377), 222aa (AY327804), 217aa (OP904044) and 194aa (OL794386) are known due to generation of TCM. The destruction of termination codon (TC) used downstream alternate termination codon (ATC) extending few AAs but generation of new TC upstream produced short length G-proteins. The 228aa G protein was produced from 219aa lineage due to TCM and used ATC. The 254aa G-protein was generated from 256aaG lineage due to early TCM (CAA=TAA; C6870T). While 261aa G-protein was generated due to TCM (TAG=CAG) and used second ATC (TAA). The generation of B-genotype of HMPV occurred due to deletion as well as extensive editing to produce 236-242aa or smaller G-proteins [224aa (OL794406), 231aa (MZ504963), 233aa (MK989732), 236aa (KC562228), 237aa (MZ504966), 238aa (KC403971), 241aa (LC769218) and 242aa (KC562242)] but B-genotypes with 219aa, 256aa or 279aa lengths G-protein were not detected. The 236aa length G-protein found in both A-genotype and B-genotype as early as 1982. The 236aa A-genotype G-protein had different C-terminal sequence than 219aaG A-genotype, 256aaG A2-genotype and 279aaG A1-genotype. Thus, we concluded 236aaG clone might be primitive HMPV lineage. We observed G-protein of A -genotype likely ended with STMQK while B-genotype 236aa G-protein ended with YNQTS. The G-protein of B1-genotype preferred HTGISPK at the carboxy-end while B2-genotype ended with SSLSS. However, mutational difference was quite high between two 219aaG or 256aaG genotypes isolated 5-10 years gap. We suggested that a new genotype method like A.236.1.US.1, A.219.1.IN.1, A2.256.1.CN.1 or A1.279.1.JP.1. This is first bioinformatics worldwide analysis of HMPV WGS sequences suggesting a new method of genotyping and compared with very neglected Indian scenario.

## Introduction

The human metapneumovirus (HMPV) is a single-stranded negative-sense human RNA virus with a genome of 13kb plus containing eight genes encoding nine proteins. Based on genomic characteristics and G-gene sequence, HMPV can be divided into four subtypes, A1, A2, B1 and B2 (Ishiguro et al. 2004; Herfst et al. 2004; Arnott et al. 2013). Further, subtype A2 was further classified into two additional subtypes A2a and A2b (Nidaira et al. 2012) while a recent classification A2c and sub sub-classification of A2b into A2b1 and A2b2 were followed (Parida et al. 2023). The major symptoms of the HMPV infection are cough, coryza (inflammation of mucous membrane of nose), chills, myalgia (muscle pain), vomiting and breathlessness. The HMPV genomic organization has close similarity (73% with 87% cover) to avian meta pneumonia virus (AMPV) while also shares close to 66% similarity at the 3’-half to the human respiratory syncytial virus (HRSV) which was shown a main virus etiology for severe pneumonia in the children of Asia and Africa (Lee et al. 2007; PERCH Study Group, 2019). The pneumonia symptoms are also found in *Streptococcus pneumoniae* bacterial infection and co-infection with HRSV. The other *Mononegavirales* like *Rhabdoviridae* rabies, the *Filoviridae* Ebola (EBOV) and Marburg (MARV), and the *Paramyxoviridae* measles (MV), mumps (MuV) and Nipah (NiV) viruses were related pathogens (van den Hoogen et al. 2002; Tiveljung-Lindell et al. 2009; Lu L et al. 2020; Gonnin L et al. 2023).

The large protein (2005aa; L protein) of HMPV has RNA-dependent RNA polymerase (RdRp) activity at the N-terminus while GDP-polyribonucleotidyltransferase and cap-specific methyltransferase activities at the C-terminus (Liang et al. 2015). The avian metapneumovirus L-protein also has close similarity to HMPV L-protein (Govindaranjan et al. 2004). *Mononegavirales* mimic RNA synthesis of their host by utilizing multifunctional RNA polymerase to replicate entire viral genomes and transcribe viral mRNAs from individual viral genes as well as synthesize 5′ methylated cap and 3′ poly(A) tail of the transcribed viral mRNAs rapidly (Liang B, 2020).

The HMPV fusion protein (F) mediates fusion of the viral envelope and cellular membranes to establish infection. The F-protein requires low pH (pH=5.2) and trypsin for fusogenic effects in Vero cells. The Asparagine 57, 172 and 353 of F-protein were glycosylated but mutation of Asparagine to Alanine did not change the fusogenic effect whereas the molecular weights of F-protein became 2kD less (Schowalter et al. 2006). Additionally, K438 could play a role potentially increasing fusion activity in association with H334 and H435 residues (Kinder et al. 2019). Due to G-gene heterogeneities, F-gene was used to consider genotyping of HMPV (Boivin et al. 2004). The very partial HMPV sequences from Canadian patients (1997-1999) with accession numbers AF371330, AF371356-67 and AF371337.2 (WGS) were used to make a phylogenetic tree in 2002 where L, F and N proteins were also attempted (van den Hoogen et al. 2001). Bastien et al and others described the heterogeneities in the G-genes of HMPV (Bastien et al. 2003; 2004; Peret TC et al. 2004).

Pinana et al disclosed first the classical 180nt duplication in the G-gene of HMPV genome (Pinana et al. 2017) and similar 111nt duplication had been shown by Saikusa et al (Saikusa et al. 2017, 2019). The 111-nucleotide duplication in the attachment gene appeared was also characterized in frequently detected genotypes (Groen K et al. 2023). Parida et al confirmed that seven out of the twenty sequences in the G-gene (accession nos. MN410888-MN410907) had 180-nucleotide duplication and eleven sequences had 111-nucleotide duplication in the A2c subtype among the Indian population (Parida P et al. 2023). But Wei et al described from WGS phylogenetic analysis that HMPV sequences with G-180nt-dup as A2b2 genotype and G-111nt-dup as A2b1genotype. The A2b1 genotype of HMPV predominates among children of Beijing, China (Wei et al. 2023). However, one drawback of the study, A2b1 clones was not compared from other countries instead A1 (KC403977), A2a (KC403979), A2b2 (KC562220) etc compared from other countries. Interestingly, we did not find 279aaG sequence class which should be A2b2 genotype suggesting a wrong interpretation of the phylogeny data!

Kikuta et al developed a rapid chromatographic immunoassay for detection of human metapneumovirus using monoclonal antibodies against nucleoprotein of hMPV (Kikuta et al. 2007). Hamelin and Boivin developed a simple enzyme-linked immunosorbent assay (ELISA) for hMPV serological testing using the nucleoprotein (N) from group A or B (Hamelin and Boivin, 2005). A complex of Phosphoprotein (P), Nucleocapsid protein (N) and L-protein (RdRP) might be occurred at the promoter during viral replication and transcription (Whitehead JD et al. 2023). The minigenome assays demonstrated that M2-2 protein inhibited both transcription and RNA replication (Kitagawa Y et al. 2009). M2-2 may also inhibit RIG-signalling after binding to RIG-1 through the inactions of CARD/TRIM25 and MAVS complexes. The inhibitory actions of HMPV M2-2 is similar to HRSV NS1 protein while similar innate immune inhibition is mediated by V-protein of few *Paramyxoviridae* viruses. The HMPV SH, a type II integral transmembrane glycoprotein, is the largest SH protein of the viral family (179aa compared to 65aa for RSV SH or 175 aa for AMPV SH). HMPV SH exists in at least three differentially glycosylated forms: un-glycosylated (23 kDa), *N*-glycosylated (>30 kDa), and highly glycosylated (>80 kDa) and may inhibit NF-κB transcriptional activity in airway epithelial cells.

During GenBank search (www.ncbi.nlm.nih.gov/nucleotide) we observed that database of recent HMPV WGS had no mention of genotyping. Such discrepancies were apparent when we multi-aligned the G-proteins of different genotypes obtained by partial sequencing, none the two G-proteins were found identical across the A1/A2 and B1/B2 genotypes. The evolutionary rate of HMPV G-gene was approximately 3.654× 10–3 substitution/site/year (95% highest probability density of 3.1303–4.0357) (Saikusa et al. 2017a,b). However, Wei et al recently collected 554 global HMPV G-gene sequences from GenBank and pointed a clear phylogenetic trait from 27 WGS like A1 (KC403977, Australia, 2003), A2a (KC403979, Australia, 2003, reported as A2-genotype in the GenBank database), A2b1 (MN745085, China, 2017, reported as A2b-genotype), A2b2 (KC562220, reported as A-genotype, 228aaG), B1-genotype (KF530163, Australia, 2004) and B2-genotype (KC562232, USA, 2001). Thus, we found a confusion in the HMPV genotype data presentation. Parida et al reported A2c lineage of HMPV from India and also have given a phylogenetic tree from PCR sequencing of G-gene (accession numbers: MN410888-MN410907) (Parida et al. 2023). The A2c genotype sequences deposited with accession numbers MN410890 and MN410893 (reported in the database as A2-genotype) with similarity to LC270124-Japan (A2b-genotype) or MF462419-China (A2-genotype) or MK947209-Croatia (no mention of genotype). They also suggested through phylogenetic analysis that KC403975-USA as A1-genotype, AY297749-Canada (no mention of genotype in the database) as A2a-genotype, KC562221-USA (reported as A-genotype) as A2b-genotype whereas AY525843-NL as B1-genotype and LC192237-Japan as B2-genotype. Similarly, Otomaru et al used different accession numbers for genotyping classifications like A1=AF371337, A2a=AB503857, A2b=AY530095, A2c=GQ153651, B1=AY525843 and B2= FJ168778 (Otomaru et al. 2023). Hence, no conclusive data obtained regarding A2a, A2b and A2c classification and mostly used partial PCR sequences with chance of 3’-end and 5’-end errors. Recently, Pragathi et al sequenced (accession numbers: OR135588-OR135607) HMPV genomes and showed the phylogenetic traits from India with PCR analysis of G-gene (Pragathi et al. 2024). We resolved the issue through the multi-alignment of G as well as N, F and L proteins but from full lengths genomic sequences. We have tried to resolve the G-protein duplication and deletions mechanisms among the different genotypes to address the horror of G-protein variations (279aa, 261aa, 256aa, 254aa, 242aa, 241aa, 237aa, 236aa, 233aa, 228aa, 219aa and 217aa etc). Such genome heterogeneities we observed in ORF7a gene of SARS-CoV-2, another 30kb positive sense RNA virus that rattled the society between 2019-2024 killing more than six million people worldwide. However, ORF7a of COVID-19 and G-protein of HMPV similarity was found minimal. The multi-alignment of G-proteins gave various length for A-genotype HMPV like 279aa (284aa), 256aa (261aa), 233aa (238aa), and 219aa (224aa) while for -B-genotype, G-proteins obtained as 242aa, 241aa, 238aa, 236aa, 237aa, 231aa, 229aa and 177aa, suggesting deletions, frameshift and termination codon mutations must be created in different HMPV genotypes during different time (1983-2024) due to immunogenic pressure in human host. Thus, we extensively reviewed the different G-genes and G-proteins here to understand the mechanisms of G-gene instability to correlate the genotype nomenclature. Surely, scientist presented genotype based on phylogenetic analysis of mostly partial genomic sequences but truly such data were not correct if you would be compared worldwide with WGS sequences. As for example, in India no full-length genome of HMPV was deposited in GenBank but PCR sequencing presented with phylogenetic analysis classifying as A1, A2, A2a, A2b (A2b1, A2b2), A2c and B1/B2 or A2.1 and A2.2.2 (Devanathan et al. 2025). BLASTN and BLASTP analysis suggested that no two G-genes or G-proteins is 100% similar in the same classification. As for example, two A2b sequences one from China (MN745086) and another from Japan (LC671555) had difference in amino acid sequence of G-protein (QGV13040 and BDE17477). We also have given the unknown genotypes of US WGS HMPV sequences between 2020-2024. The few corrupted HMPV sequences are also described that may cause problems during multi-alignment and phylogenetic analysis. Surely, we pinpointed and compared the molecular basis of different genotypes based on G protein through extensive WGS multi-alignment (CLUSTAL-Omega) and BLASTP as well aa BLASTN analyses. We solved the problem of genotyping by simple assumptions that 219aaG clone is primitive termed as A-genotype from where 180nt duplication had occurred giving A1-genotype and further deletion occurred to produce most recent 256aaG clone A2-genotype. The B-genotype clones was derived from 236aa B-genotype whereas B1 and B2 genotypes had clear divergence in amino acids in the G-protein and also had difference at the C-terminal ends.

## Method

We used CLUSTAL-Omega software for multi-alignment of DNA and protein sequences (Corpet, 1988; Sievers et al. 2011). The MultAlin software was also used to check amino acid similarity with color presentation. Sequence similarity between the two sequence was determined by BLAST2 Program. However, BLAST2 search between A-genotype and B-genotype may give incomplete data due to fragmentation near insertion point and in such cases two proteins were aligned by CLUSTAL-Omega program. Extensive search of HMPV genotypes was made at www.ncbi.nlm.nih.gov/protein/nucleotide. SWISS-Modelling was used to predict the structure of G-protein (Waterhouse et al. 2018)

The penetration of specific mutated HMPV peptide sequences were performed by NCBI BLASTP search using 60aa of G-protein (Lu et al. 2024). When we found an extended G-protein with amino acid difference due to frameshift, we immediately did BLASTP search to check the penetration of such peptide in the database. In many cases, only one sequence appeared as 100% identity and we presumed that such data might be a sequencing error. We like greater than 99% similarity between two sequences of the same genotype but near 88% RNA sequence similarity was found between A-genotype and B-genotype. Thus, presently, A1/A2 and B1/B2 genotype classification could be made even a strong bias of 180nt and 111nt duplication mechanisms in the G-gene. If A2b1 classification is 180nt duplication and A2b2 for 111nt, then a strong bias between A2c classification for Indian strains seems problematic (Parida et al. 2023). We found very neglected genotyping throughout our investigation. Thus, we tried to resolve the issue by checking the important single amino acid point mutation between two genotypes or sub-genotypes introducing country name in the genotype suffix.

Various diagnostic tools were designed to detect the infectious agents. The FilmArray Respiratory Panel (RP/RP2) (BioFire Diagnostics, Salt Lake City, Utah, USA) multiplex PCR was used to detect 19 organisms including adenovirus, coronavirus (229E, HKU1, NL63, OC43), human metapneumovirus, human rhino/enterovirus, influenza A H1, influenza A H1 2009, influenza A H3, influenza B, Middle Eastern respiratory syndrome coronavirus (MERS-CoV), parainfluenza serotypes 1–4, respiratory syncytial virus (RSV) among viruses and *Bordetella pertussis*, *Bordetella parapertussis*, *Chlamydia pneumoniae* and *Mycoplasma pneumoniae* among bacteria (Maertzdorf et al. 2004; Sugimoto et al. 2023). The HMPV was detected in 2% samples during August-November between Monsoon and Winter seasons in a south Indian study (Anand M et al. 2020). However, influenza-A virus infection was found high as compared to HMPV and HRSV. Parida et al described the method of sequencing of 180nt and 111nt duplication in the A2c genotype of HMPV using Superscript III Platinum™ One-Step qRT-PCR Kit (catalogue. No. 1821198, Invitrogen) and further sequenced using BigDye Terminator (version 3.1) cycle sequencing kit (Applied Biosystems) according to the manufacturer’s protocol. Purified sequences were analysed using a 3500xL Genetic analyser (Applied Biosystems). The data indicated accession numbers MK947209, LC192241, MF462419 aligned between A2c genotype sequences like accession numbers MN410905 and MN410900 at the one side and MN410890 and MN410895 at the other side of the phylogeny. We used such sequences to coin the A2C genotype WGS from other countries by BLAST search similarity. The full length A2c sequence of HMPV was not reported from India as well as from other countries (Chaudhary et al. 2014)!

## Result

### Analysis of full length HMPV genomes of different genotypes to check the duplication and deletions

The multi-alignment and phylogram of WGS genomes of HMPV was demonstrated in figure-1A and figure-1B. The amino acid lengths of G-proteins are shown with genotype and country. The phylogram indicated a clear division between A- and B-genotype as well as clarity of B1 vs B2 genotypes were seen. The G-protein amino acid lengths clearly contributed to the phylogram as shown in figure-1B. Thus, phylogenetic analysis of 219aaG, 279aaG and 256aaG lineages of complete HMPV genomes and their classification into A-genotype (219aaG), A1-genotype (279aaG) and A2-genotype (256aaG) was justified. The hypothesis was that 219aaG clone duplicated 180nt to form 279aaG clone which subsequently lost 69nt by deletion forming the most recent 256aaG clones. The point mutations in primitive 236aaG variants disclosed all A, A1 and A2 genotypes between 1982-2010 like due to difference in few point mutations. However, clear spread of 256aaG lineages with more point mutations made it difficult to use same genotype classification as point mutations strong varied. Since then, scientists did not disclose the genotype of HMPV during GenBank submission and although B-genotype sub-classification (B1 and B2) appeared normal.

**Fig. 1A.**
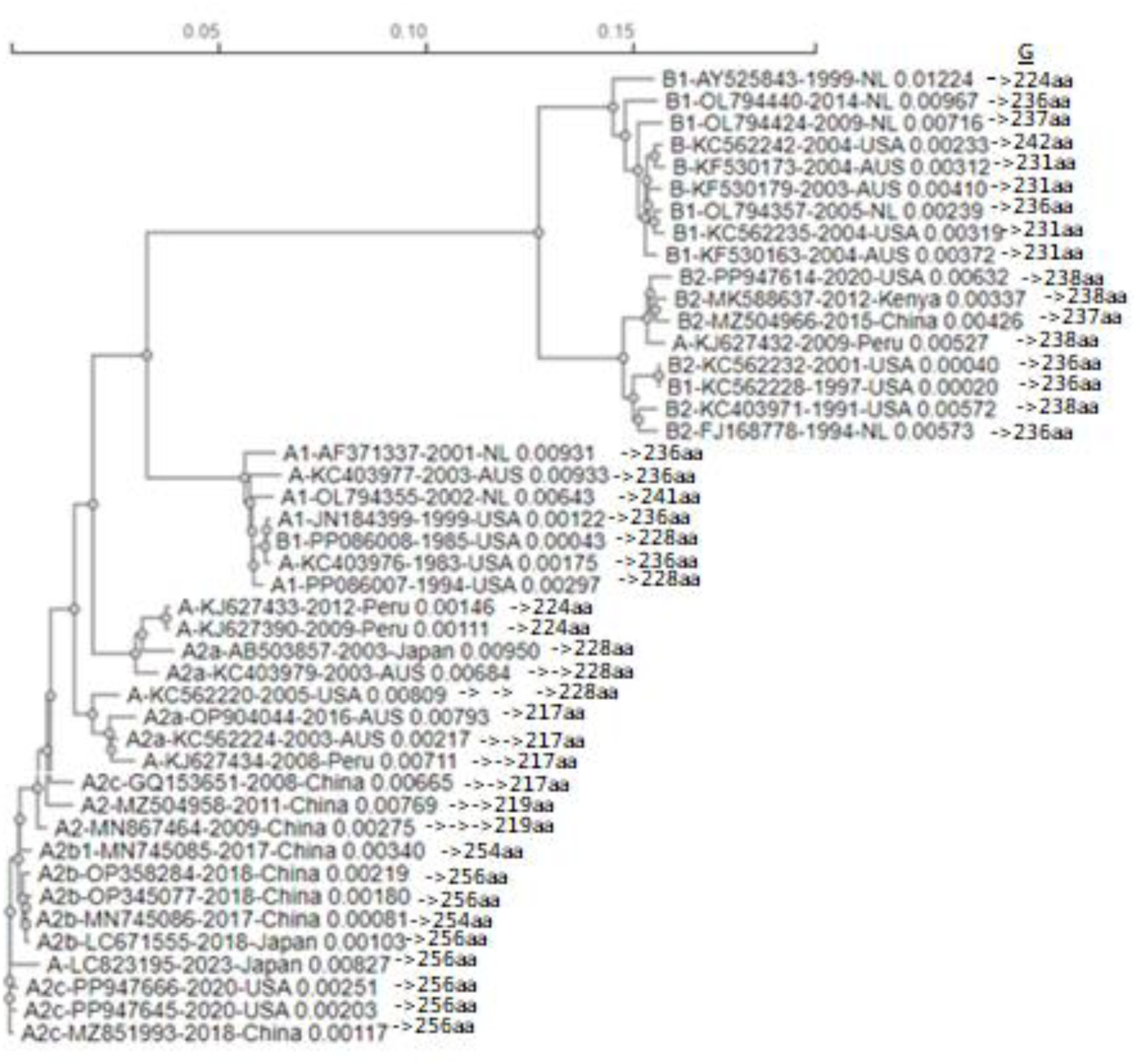
The multi-alignment and phylogeny of WGS genomes of HMPV. The amino acid lengths of G-proteins are shown with genotype and country. The 279aaG clone was not used here. Data indicated a clear division between A- and B-genotype as well as clarity of B1 vs B2 genotypes are there. The G-protein amino acid lengths clearly contribute also to the phylogeny. It should be noted that KJ627432 and PP086008 had error in genotyping. The A2C genotype WGS sequences were compared with partial sequences from India (Parida P et al. 2023)

**Fig. 1B.**
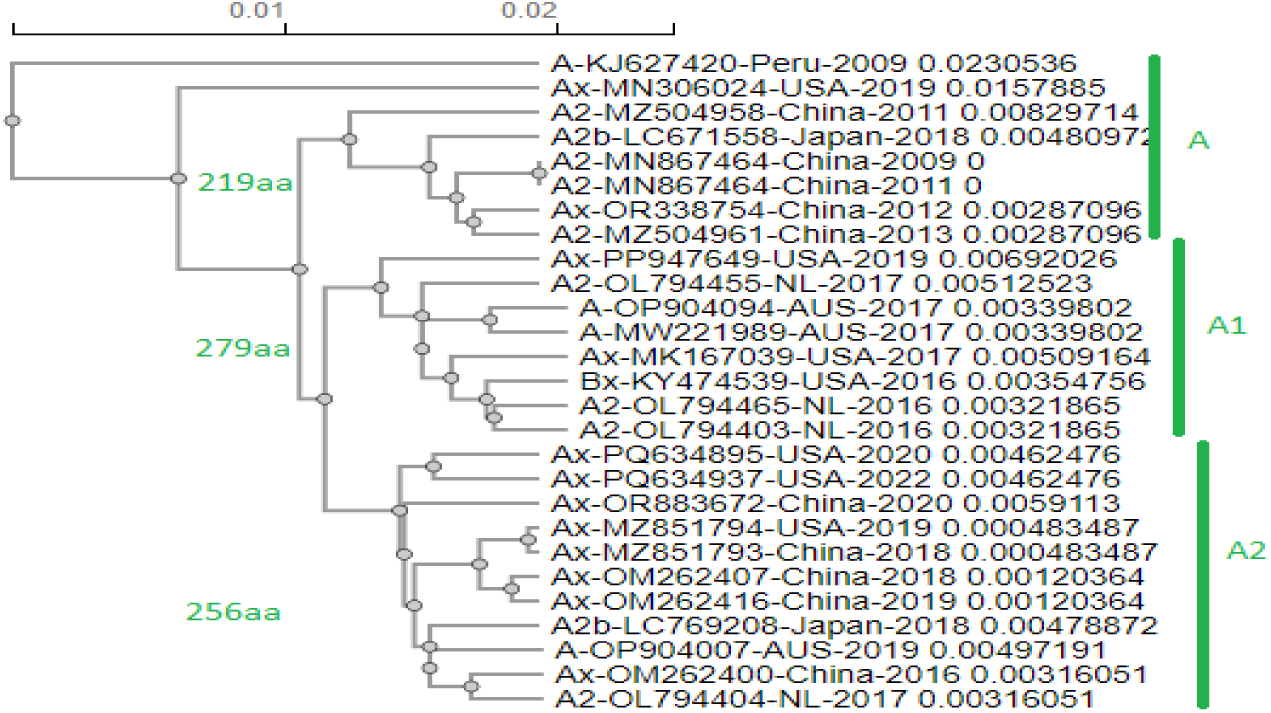
Multi-alignment and Phylogenetic analysis of 219aaG, 279aaG and 256aaG lineages of complete HMPV genomes and their classification into A-genotype (219aaG), A1-genotype (279aaG) and A2-genotype (256aaG). The hypothesis was that 219aaG clone duplicated 180nt to form 279aaG clone which subsequently lost 69nt by deletion forming the most recent 256aaG clones. The phylogeny proved optimistic with the length of the G-protein even WGS of HMPV used for multi-alignment. Ax means the A-genotype and Bx means B-genotype as judged by multi-alignment with known A-genotype and B-genotype sequences. Many scientists did not disclose the genotype of HMPV during GenBank submission.

The multi-alignment of few complete HMPV sequences was shown in figure-2A to demonstrate the duplication and deletion in the genomes. Similarly, BLASTN search of the deleted region of the genome as in figure-2B demonstrated a 111nt duplication (we called 69nt deletion from 279aaG clone) in the WGS of HMPV genome. The genome size of HMPV varies from 11300-13450 nucleotides for A-genotype and 11250-13000 for B-genotype suggesting there was various deletions in the genomes but we found more deletions in B-genotype lineages (Piyaratna et al. 2011). We used the WGS clones with intact N-protein at the 5’-end and L-protein at the 3’-end but 3’ and 5’ UTR sequences might be different. The multi-alignment search with different G-proteins always demonstrated the duplication amino acid regions as demonstrated in figure-3A (279aaG) and figure-3B (256aaG). Then, we tried to classify the deletions on the basis of A/B genotypes but it was hard as complete genome of A2a and A2c genotypes were not found and those of A2b genotype was available in the database. Thus, we proposed a model where all 219aaG as well as 236aaG were A-genotype, 279aaG are A1-genotype and 256aaG as A2-genotype. We put the accession numbers of HMPV in table-1 according to the date of virus isolation, genotype, genome length, G-protein length to get access of WGS for further analysis and comparison. One of the observations was that 60aa N-terminal amino acids of G-proteins were comparatively less mutation prone but partial sequences of G-protein was a problem in multi-alignment. Surely, PCR sequencing at the both terminuses are likely error prone. So, we BLASTP search of G-protein 61-120aa to find different population of genotypes based on G-protein sequence as well as to compare the WGS available by multi-alignment. However, G-protein from complete HMPV sequences gave a clear picture of B1 and B2 genotype as well as A-genotypes. We explained the basis of various lengths for A-genotype HMPV G-proteins like 279aa (284aa), 256aa (261aa), 233aa (238aa), 219aa (224aa) and 217aa (222aa). In sate values are G-protein lengths when upstream second initiation codon was considered and such method was reported by Netherland scientists in the database.

**Fig. 2A.**
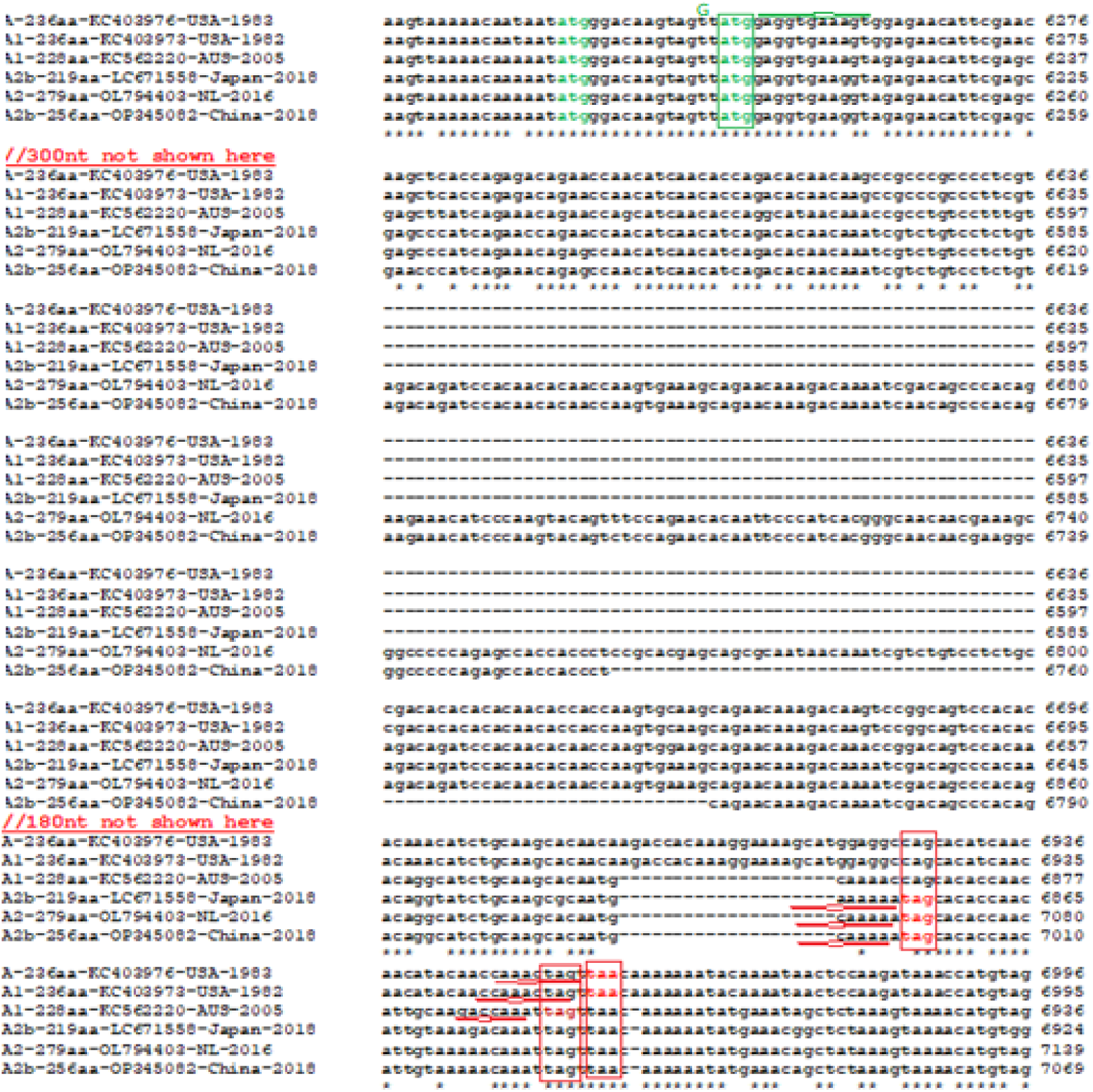
The multi-alignment of different human meta pneumonia virus (HMPV) A-genotype genomes to check 180nt and 111nt duplication in the G-gene. The 229aaG clone converted into 279aaG clone by 180nt duplication which had converted into 256aaG clone by 69nt deletion. The 236aaG clone also had 180nt deletion but insertion near termination codon and using alternate termination codon. The 228aaG clone also had 180nt deletion but a deletion near termination codon, Such, 20nt deletion also appeared in the 256aaG clone and 279aaG clone suggesting 236aaG clone was appeared early. As the 236aaG clone appeared first (as early as data deposited between 1982-1983), it was likely the 219aaG clone appeared from 236aaG clone during the early generation and transmission of HMPV.

**Fig. 2B.**
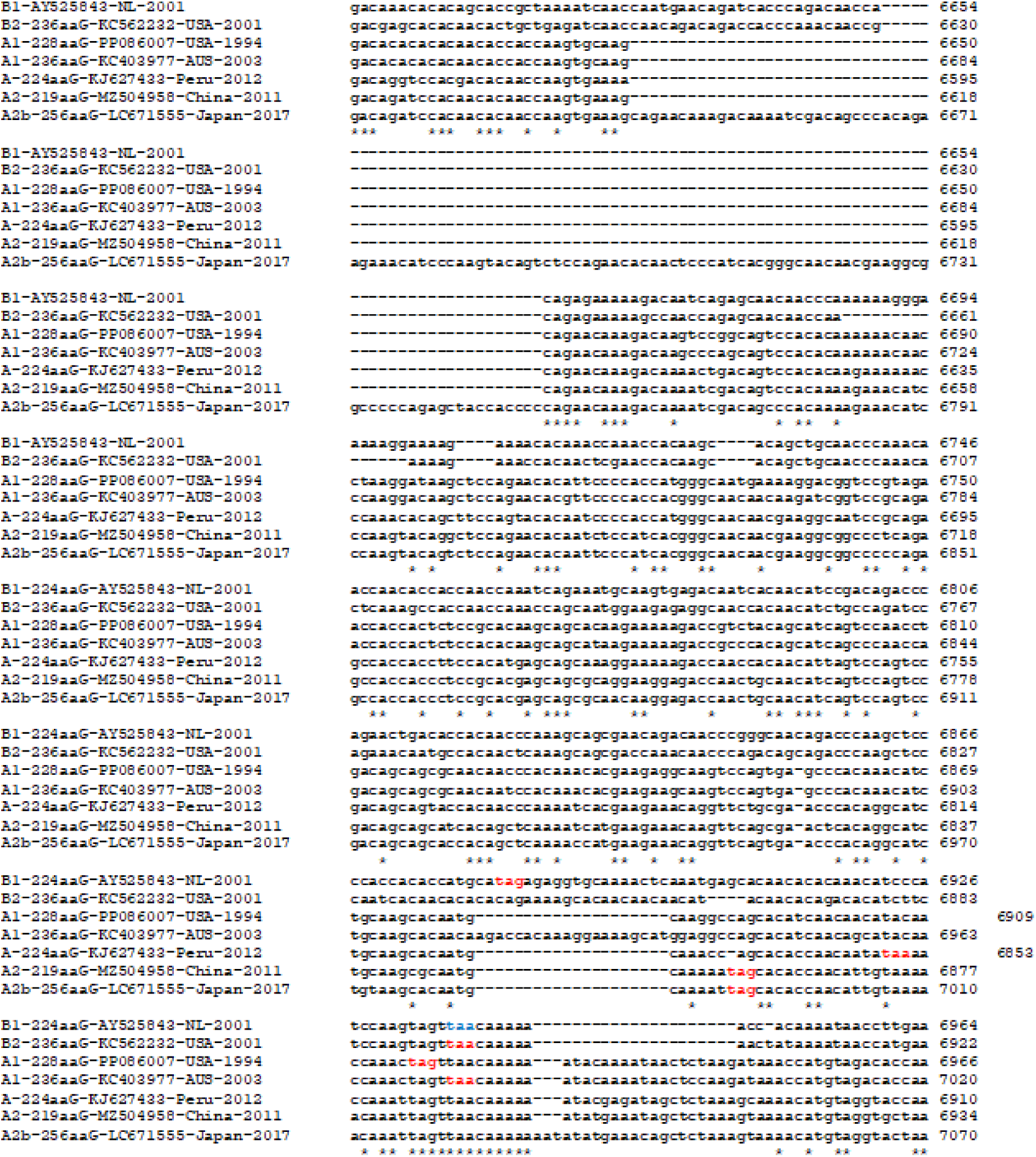
Multi-alignment of HMPV G-gene region of A-as well as B-genotypes to demonstrate 111nt duplication in 256aaG A2-genotype clone (LC671555). The 224aaG B-genotype clone (KC562232) was produced from 236aaG B-genotype due to upstream TCM (AY525843) while 224aa A-genotype clone (KJ627433) might be produced due to one amino acid deletion in the termination codon and used downstream termination codon extending few amino acids and likely was generated from 219aaG clone (MZ504958). However, sequencing error in Peru clone (KJ627433) possible but such deletion was confirmed in hundred clones worldwide. Truly, it is very hard to align between A- and B-genotypes genomes due to different lengths of G-proteins. Termination codons are shown red.

**Fig. 3A.**
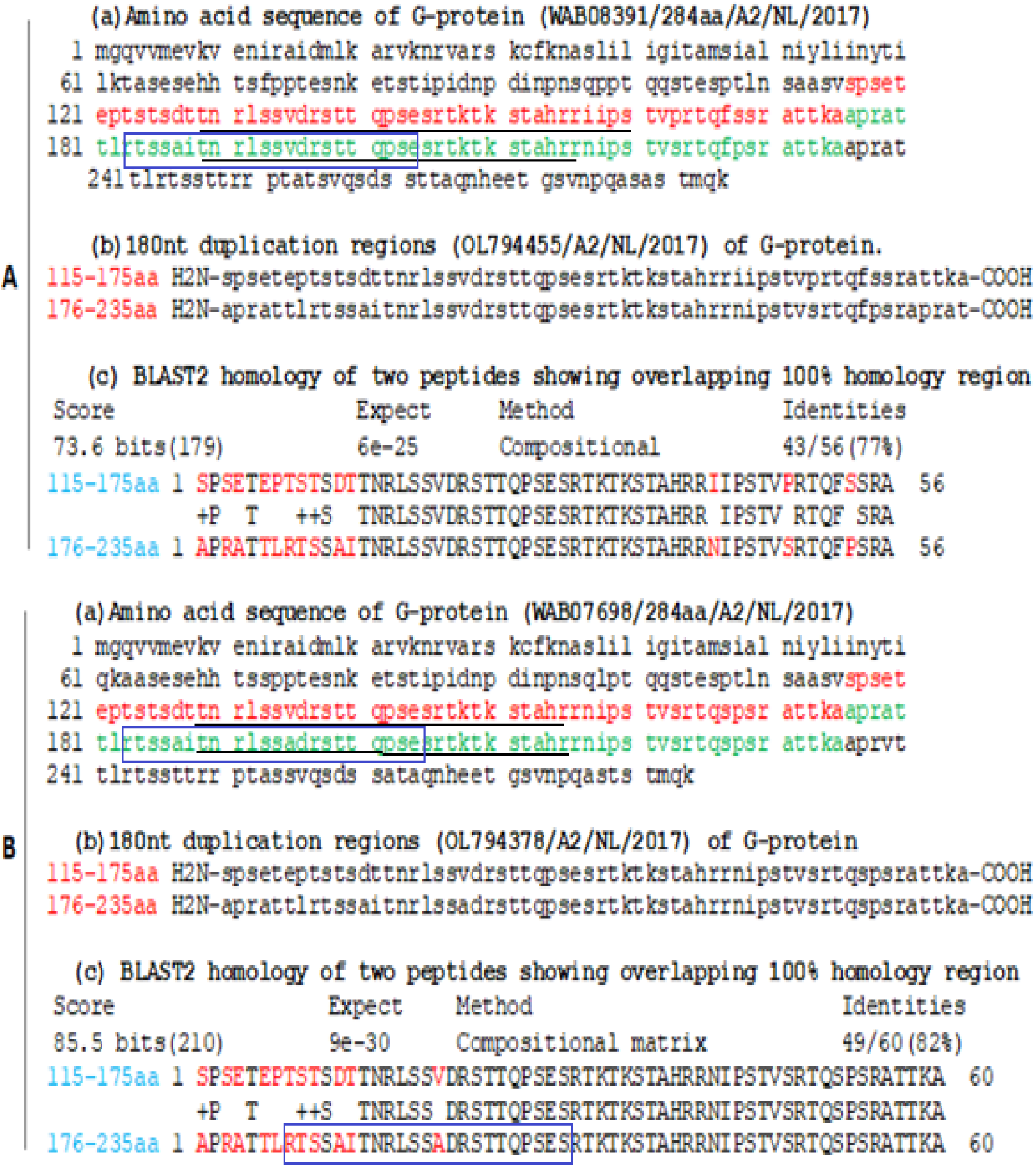
Demonstration of 60aa duplication (180nt) in the G-proteins (WAB08391 and WAB07698) of two Netherland HMPV A2-genotype lineages (OL794455 and OL794378). No homologies at the NH2-terminus 13aa but 76% and 81% homologies were detected with few point mutations respectively. The role of 279aa G-protein repetitive amino acid sequences in virus clinical manifestation was not clear. The 3-D structure of different G-proteins were not investigated yet!

**Fig. 3B.**
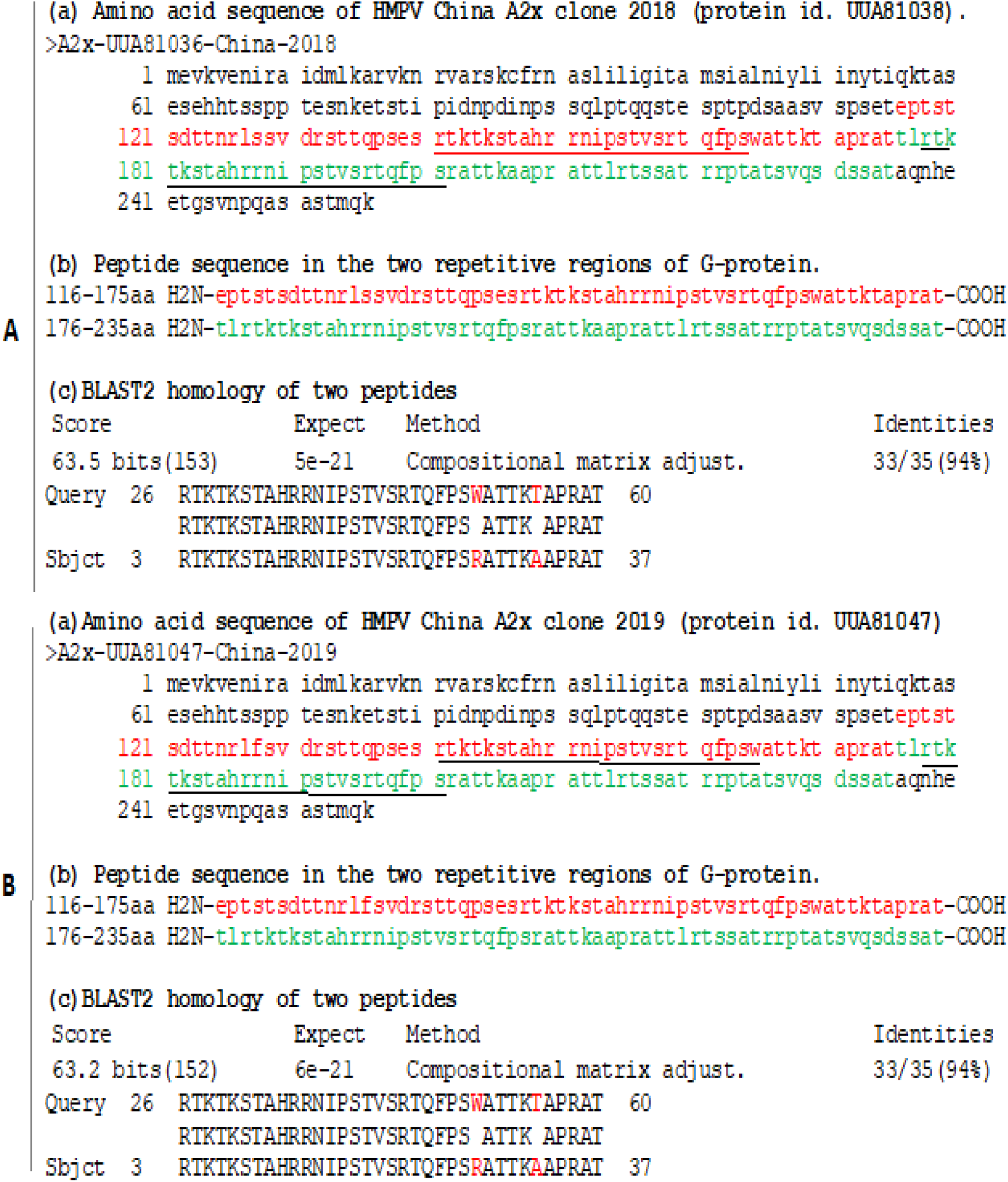
Demonstration of 111nt duplication and amino acid changes in the G-proteins (UUA81038 and UUA81047) of two China HMPV A2x-genotype lineages (MZ851793 and MZ851794). No homologies at the NH2-terminus 25aa but 94% homologies were detected with few point mutations at the Carboxy terminus. The 111nt duplication appeared due to deletion of 69nt from 180nt duplication 279aaG lineages. We surprisingly found both 279aa and 256aa A2x clones had repetitive sequences with ∼46aa homology and ∼36aa homology respectively. The H2N-RTS SAI TNR LSS VDR STT QPS ES-CO2H amino acid sequence was absent in G-protein of 279aa (284aa) clone as compared to 256aa clone and thus, the deletion was occurred in the 2^nd^ repetitive sequence of the 279aaG clone (blue box). The amino acid positions were shown. (a) amino acid sequence of 256aaG (b) two repetitive G-sequences in the same protein and (c) BLASTP search produced the exact homology region in the same G-protein. The consequence of such repetition was not known in term of 3-D structure, immunogenicity and attachment to host cells.

In figure-4, we demonstrated the multi-alignment of different G-proteins of A-genotype group. We found the 219aaG, 256aaG and 279aaG preferred H_2_N-SASTMQK-CO_2_H amino acid sequence at the end. It appeared 236aaG protein was produced by destruction of known termination codon and used downstream termination codon extending new 18aa sequence at the carboxy-terminus and ended with H_2_N-TTYQNTS-CO_2_H.

**Fig. 4.**
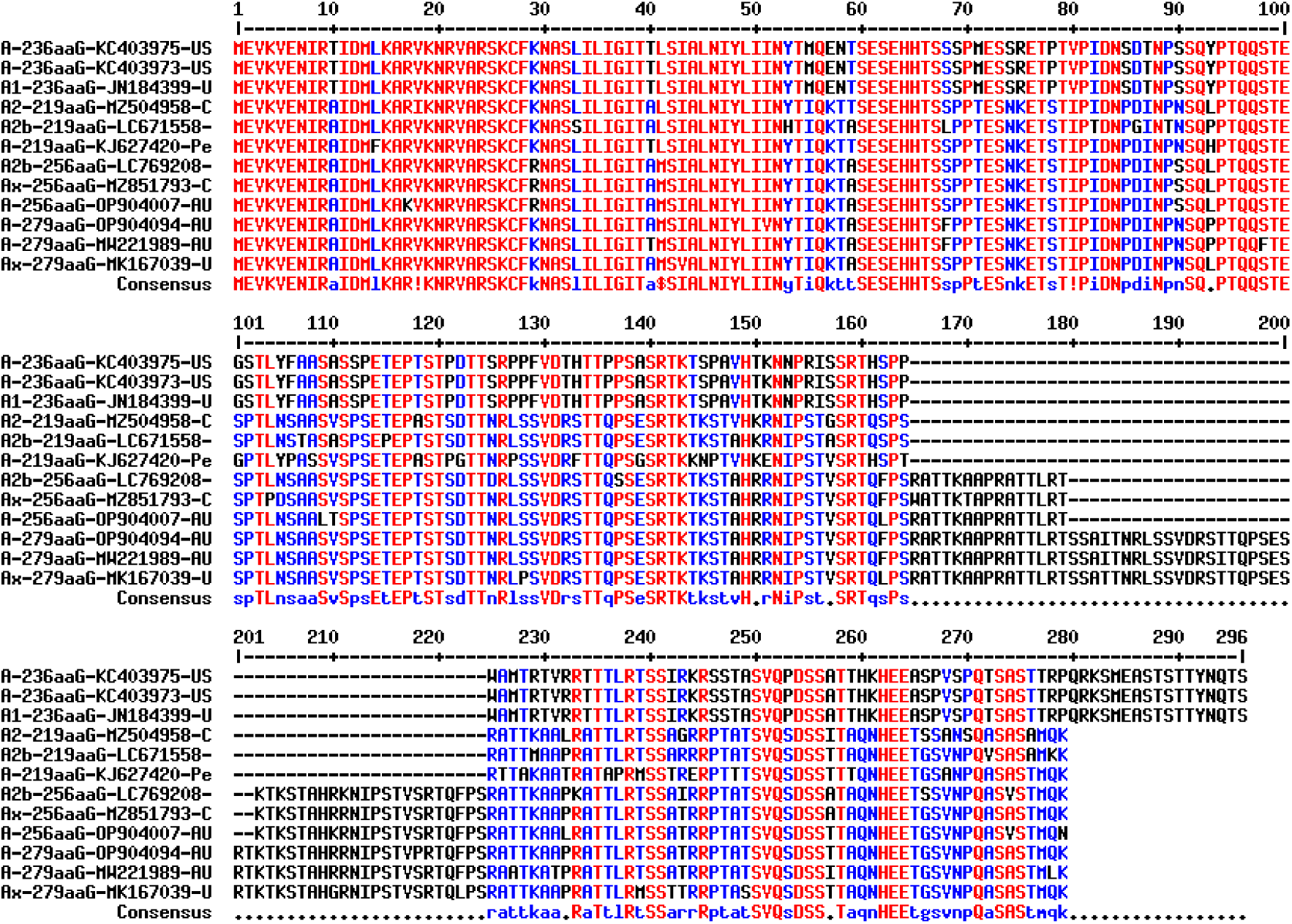
Multi-alignment of different lengths of A-genotype G-proteins (219aa, 236aa, 256aa and 279aa) to demonstrate 236aa G-protein had no carboxy-terminal homology with other three partners. The synergy in sequence during duplication and deletion were observed for 219aa, 256aa and 279aa G-proteins. It appeared 236G protein was produced by destruction of known termination codon and used downstream termination codon extending new 18aa sequence at the carboxy-terminus.

**Table 1:**
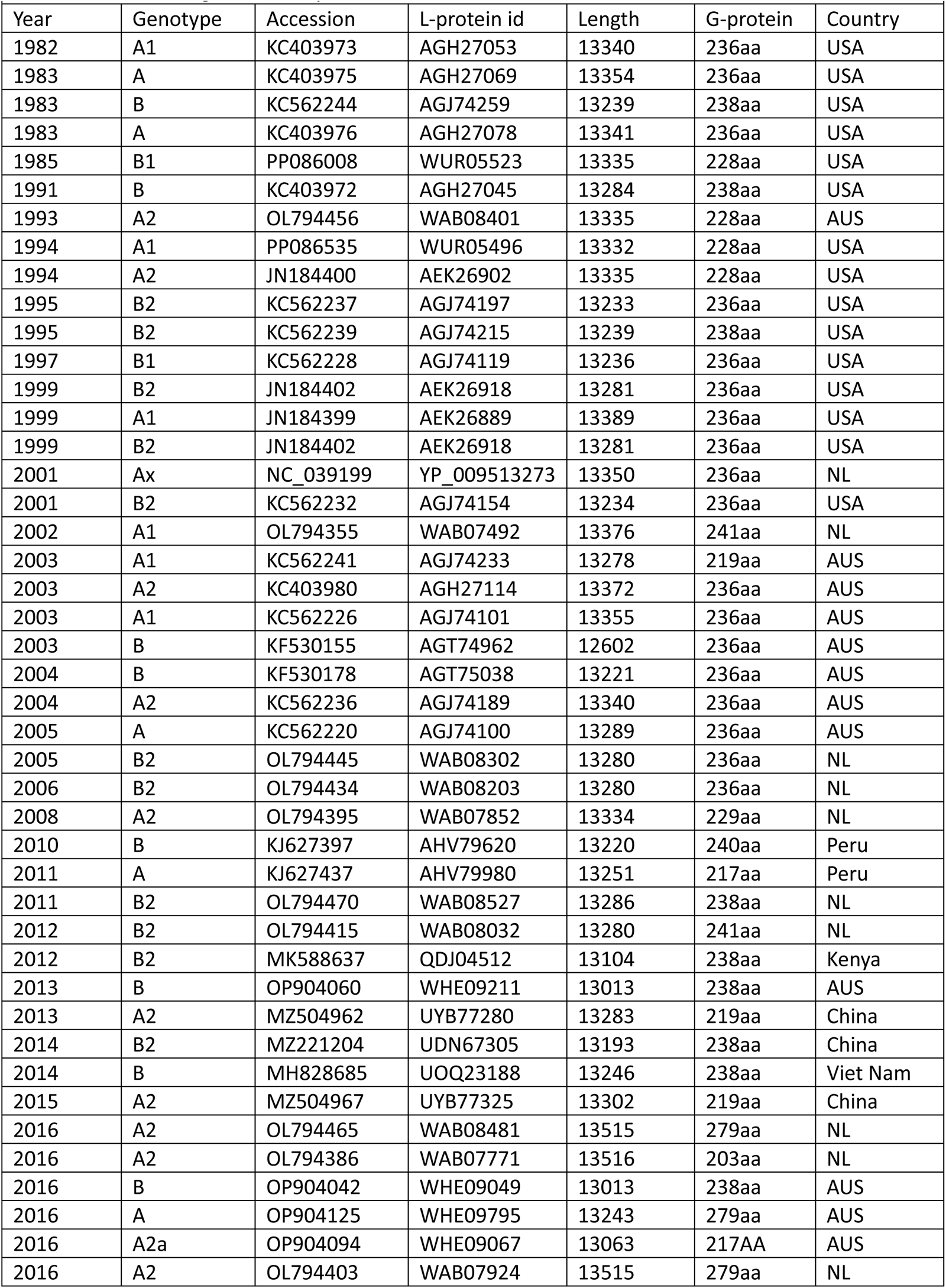
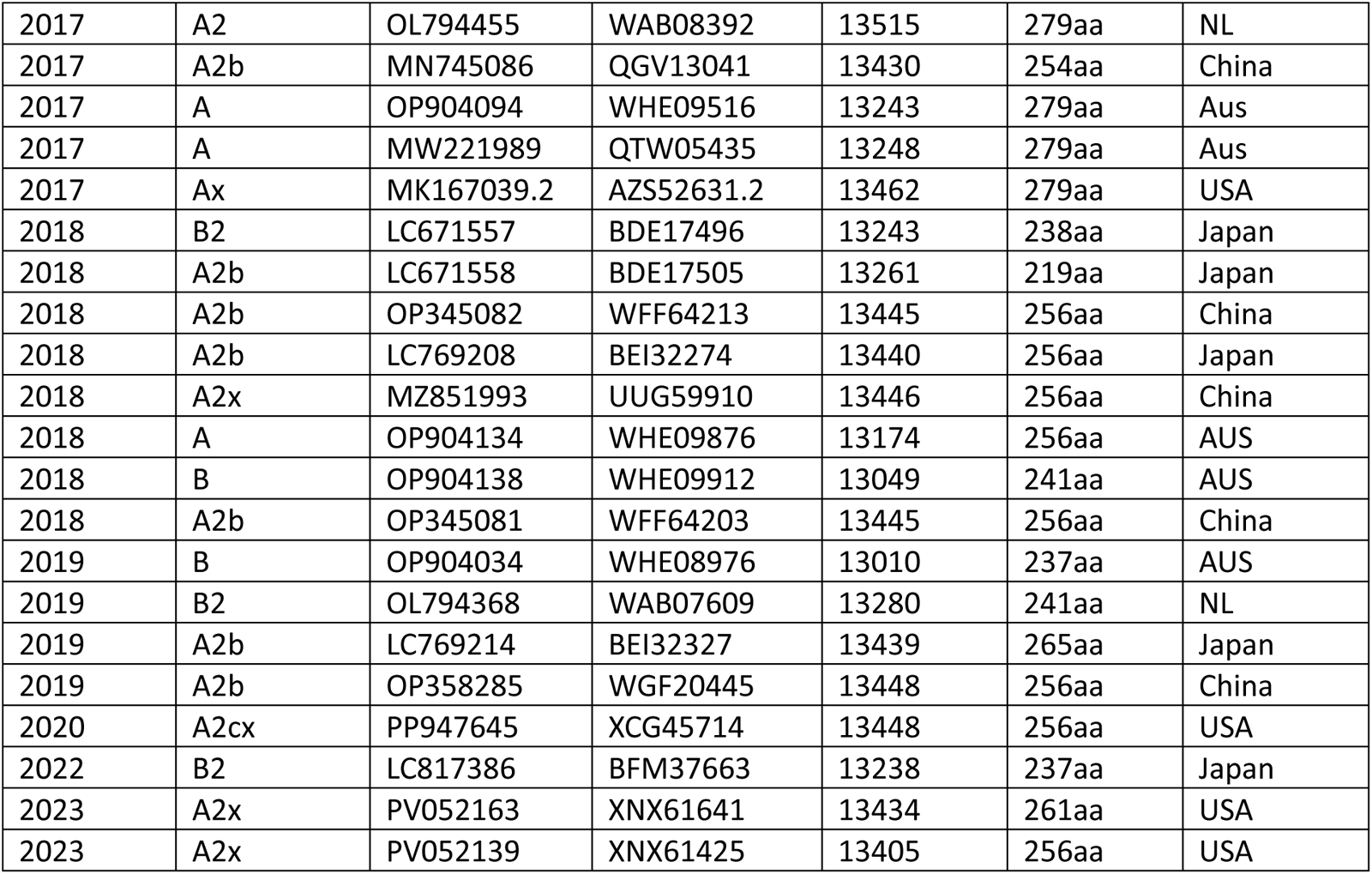
Characteristics of few important full length human metapneumovirus insertion and deletion mutants for multi-alignment analysis and BLAST search.

Multi-alignment of G-proteins of different HMPV B-genotypes were also performed (figure-5). Sadly, the most US HMPV clones from 2015-2025 did not reported genotype classification. We denoted by Ax and Bx after alignment with known genotype with 98-99% homology. It appeared that 240aaG clones had gone minor deletions to produce 238aaG (161EK or 161GK deletion), 237aaG (EKE or GKE or EKD or GKD) and 236aaG (GKEK or EKER or GKDK or EKDK) clones. B1-genotype ends with H_2_N-HTGI(S/L)(P/S)K-CO_2_H and B2-genotype ends with H_2_N-DTSS(L/P)SS-CO_2_H. The alteration of termination codon due to point mutation might cause pre-termination ending with H_2_N-SAGPRRT-CO_2_H and H_2_N-(S/G)AGPR-CO_2_H. We also established that 64L=B-genotype, 64N=B1-genotype and 64H=B2-genotype. We also found few B1/B2 specific amino acids: 97W, 114Y, 161R and 190S for B1-genotype while 97L, 114H, 161K and 190R for B2-genotype. G-protein length 231aa and 241aa belong to B1-genotype whereas 236aa, 237aa, 238aa and 240aa belong to B2-genotype. The exceptional genetic variants are also there due to frame shift as we saw in G-protein with protein id XDK70484. The AAS92886 224aa protein also terminated early ending with H_2_N-SSPPHHA-CO_2_H while in 207aa clone (ABC26383) ended far early with H_2_N-SSOQTT-CO_2_H (figure-5). We did not find HMPV B-genotype clone with amino acid lengths 219 (224), 256 (261) and 279 (284) which exclusively for A-genotype. Thus, we accepted that 219aa changed to 279aa due to 180nt duplication (A1-genotype) and then 69nt deletion produced 256aa (A2-genotype). Then, further recombination and extensive editing produced B-genotype with 240aa or 236aa. We have no clue for the absence of high molecular weight G-protein (256aaG and 278aaG) for B-genotype (Yang et al. 2013)!

**Fig. 5.**
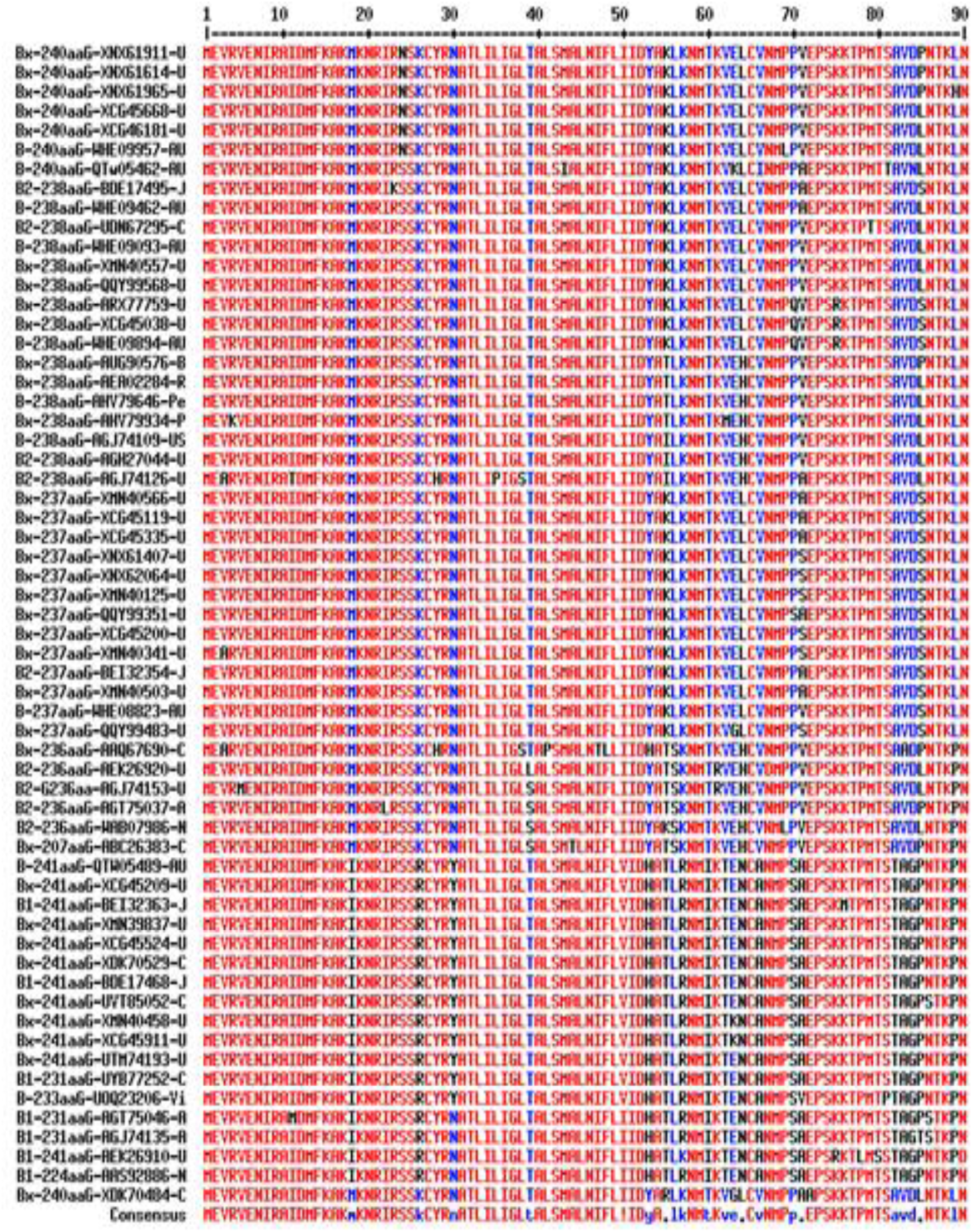
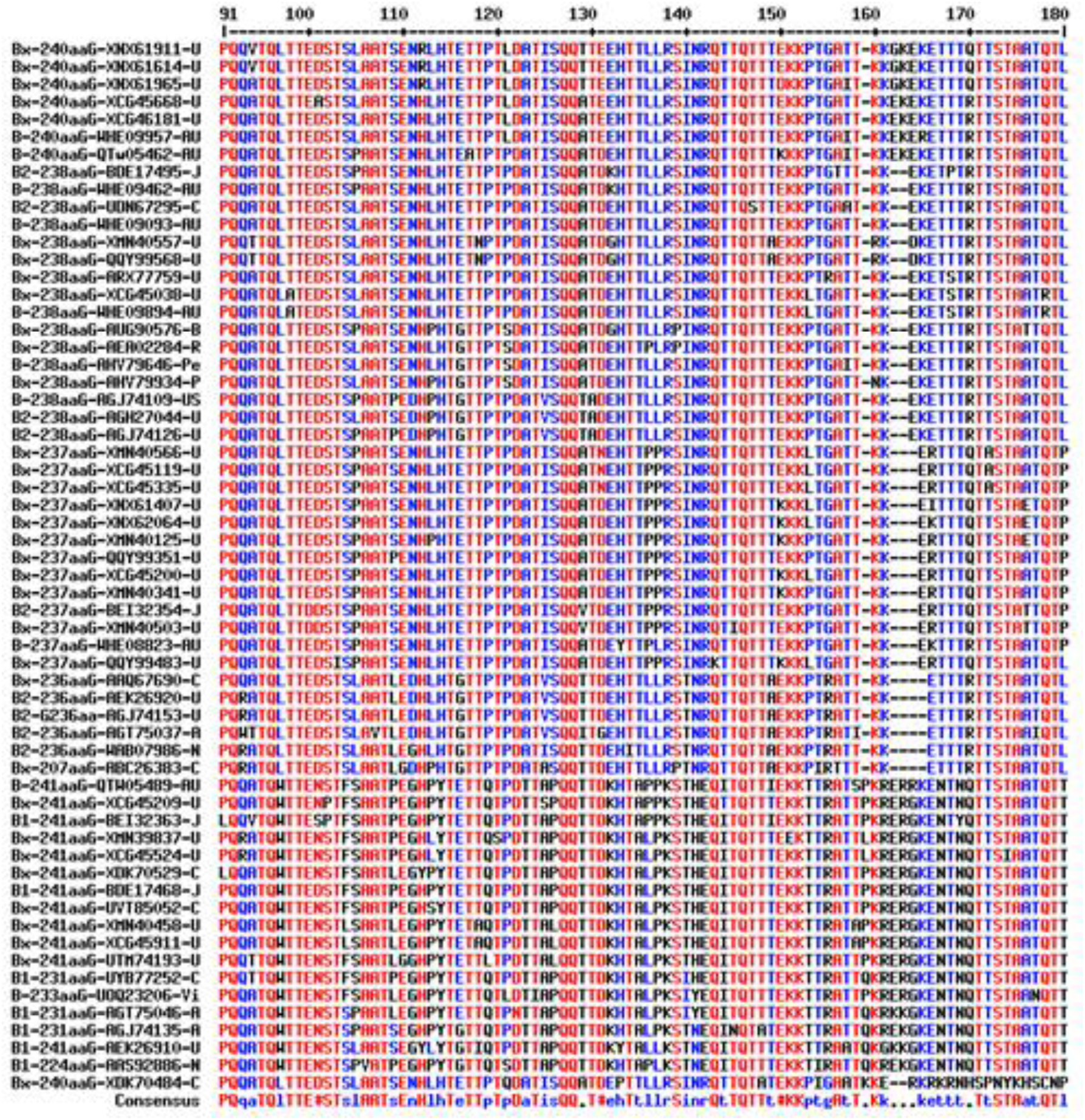
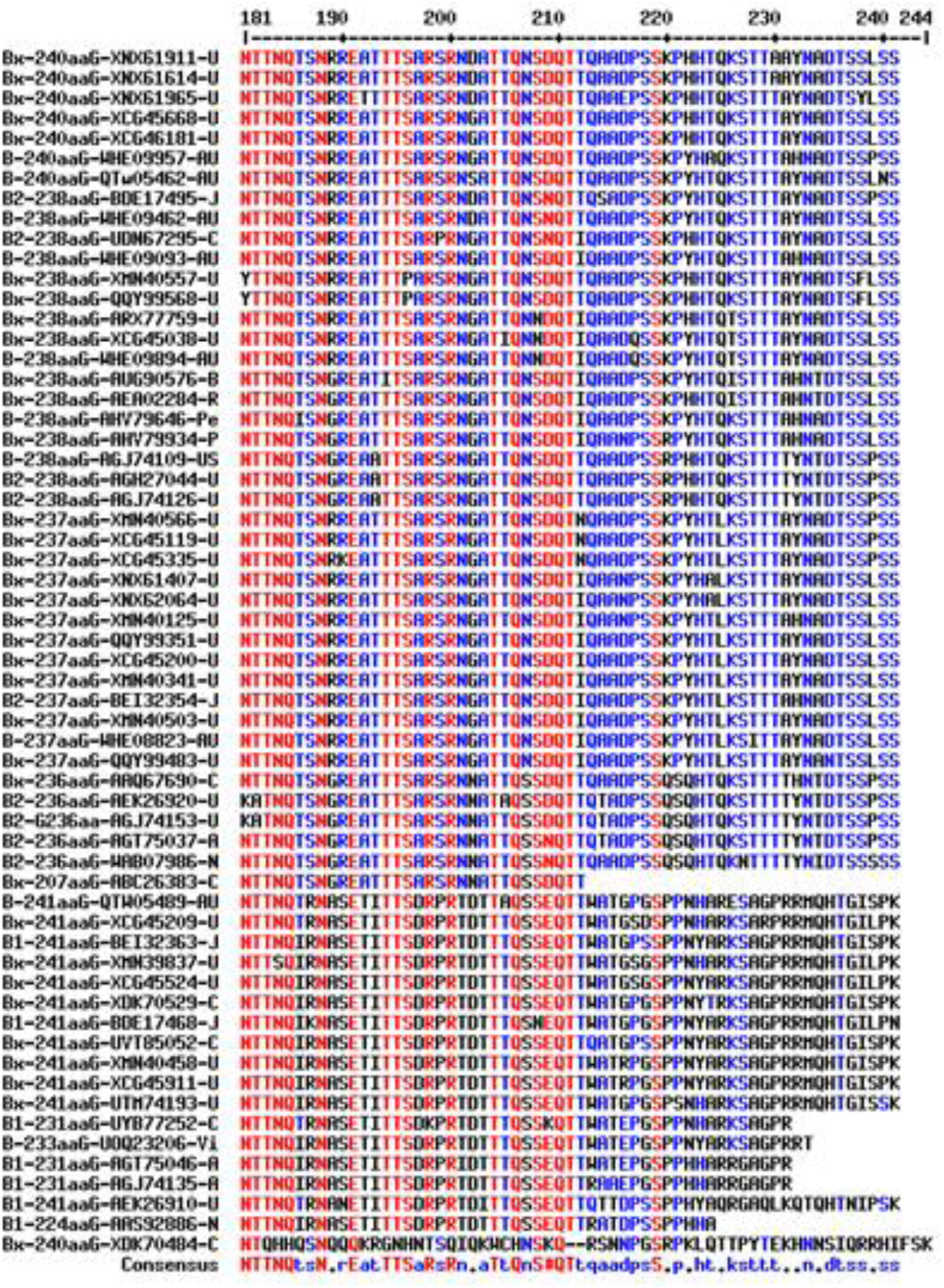
Multi-alignment of G-proteins of different HMPV B-genotypes. The genotypes of US clones were not mentioned in the database and denoted by Bx. It appeared that 240aaG clones had gone minor deletions to produce 238aaG (161EK or 161GK deletion), 237aaG (EKE or GKE or EKD or GKD) and 236aaG (GKEK or EKER or GKDK or EKDK) clones. B1-genotype ends with HTGI(S/L)(P/S)K and B2-genotype ends with DTSS(L/P)SS. The alteration of termination codon due to point mutation might cause pre termination ending with SAGPRRT and (S/G)AGPR. We also established that 64L=B-genotype, 64N=B1-genotype and 64H=B2-genotype. We also found few B1/B2 specific amino acids: 97W, 114Y, 161R and 190S for B1-genotype while 97L, 114H, 161K and 190R for B2-genotype. G-protein length 231aa and 241aa belong to B1-genotype whereas 236aa, 237aa, 238aa and 240aa belong to B2-genotype. The exceptional genetic variants are also there due to frame shift as we saw in XDK70484. The AAS92886 224aa protein also terminated early ending with SSPPHHA while in 207aa clone (ABC26383) ended far early with SSOQTT. We did not find HMPV B-genotype clone with amino acid lengths 219(224), 256(261) and 279(284) which exclusively for A-genotype. Thus, we accepted that 219aa changed to 279aa due to 180nt duplication (A1-genotype) and then 69nt deletion produced 256aa (A2-genotype). Then further recombination and extensive editing produced B-genotype with 240aa or 236aa.

**Fig. 6A.**
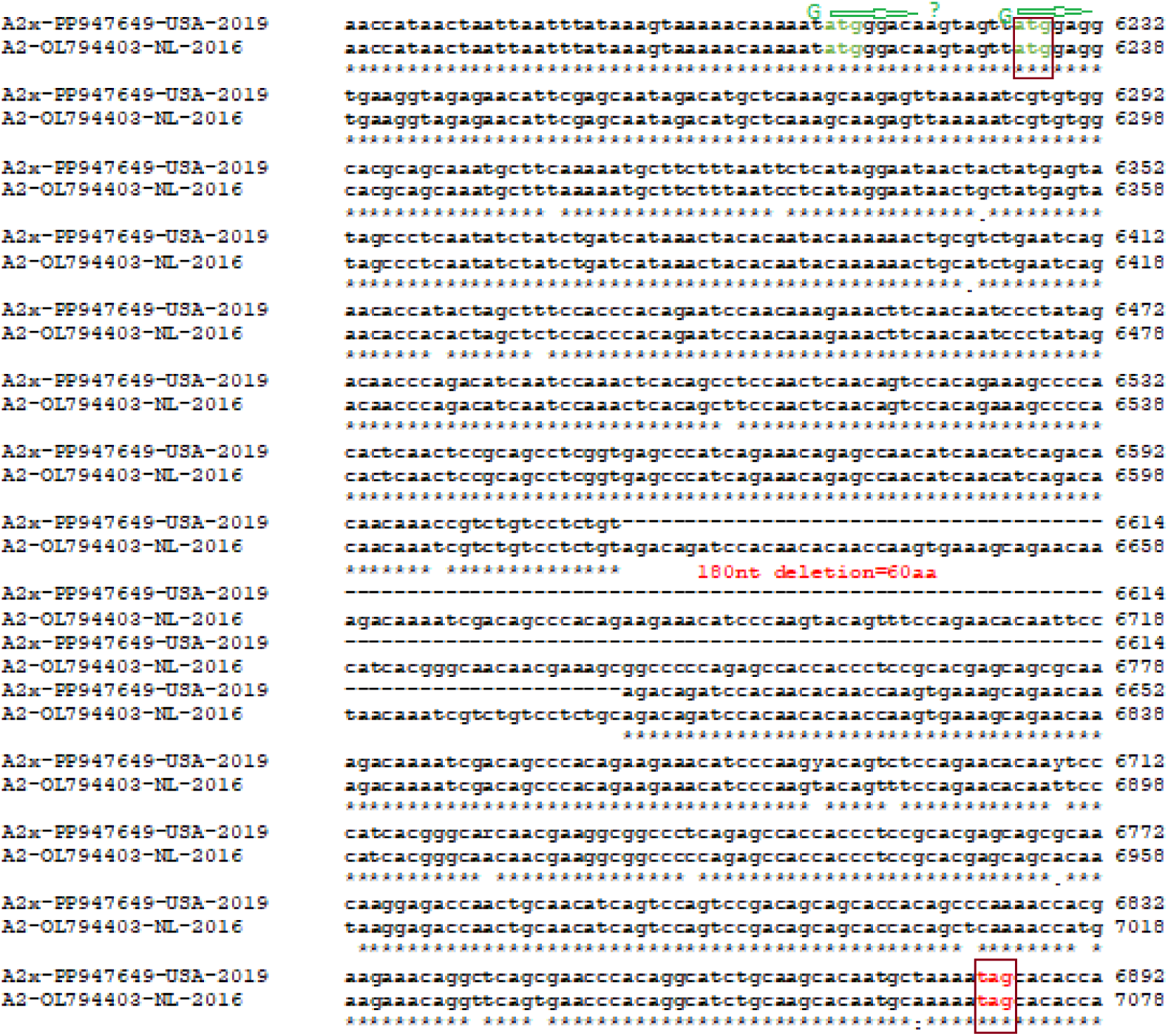
Example of in reading frame 180nt insertion into G-gene forming 279aa instead early A-genotype 219aa G-protein of HMPV (OL794403). The genotype of US-clone was not reported but similar to A2-genotype.

**Fig. 6B.**
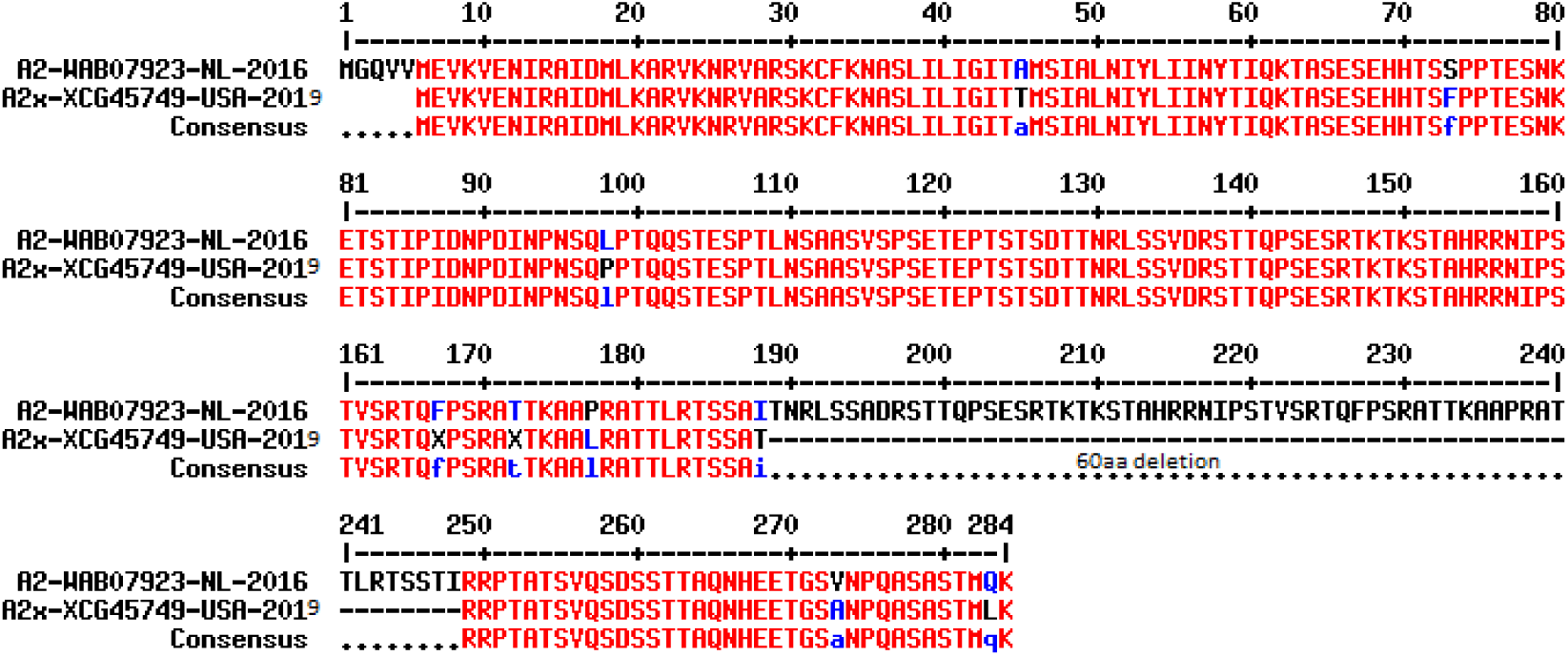
The G-protein amino acid differences (WAB07923=284aa; XCG45749=219aa) between two A2 genotypes of HMPV (PP947649 and OL794403) as shown in figure-6A due to 180nt duplication in the middle but remained in reading frame. We also found seven point-mutations whereas the 162 and 167 amino acids were not mentioned in US-clone. The upstream G-protein ATG initiation codon was proposed in the Netherland-clone making 284aaG..

### Characterization of termination codon mutation making varying lengths of G-proteins

At this point, we confirmed the various lengths G-proteins of HMPV of both A-genotype and B-genotype. We have tried next to find the mechanism of TCM due to point mutation, deletion, recombination and most accepted mechanism, duplication (Zhong et al. 2012). We showed the generation of 180nt and 111nt duplications in the G-gene of HMPV as demonstrated in figure-6 and figure-7 where 219aaG converted into 279aaG and then converted into 256aaG lineages. In figure-8, we demonstrated the mechanism of HMPV 219aaG (224aaG) G-gene changed into 228aaG (233aaG) clone due to point mutation (6889T>C) destroying termination codon (TAG=CAG; OL794397) and extended 228aa G-protein (protein id. WAB07869) was produced from the second TAG termination codon. The alignment of 236aaG WGS (KC403975) with 228aaG WGS (KJ627377) produced a 20nt gap near the termination codon (TAG, same both cases) suggesting 228aaG clone was derived from 219aaG clone. Further, the greater homology detected between two G-proteins (AGH27068, 236aa and AHV79442, 228aa) at the carboxy terminus short extension. If 228aaG clone produced from 236aaG clone then about fifty amino acids would be changed due to frame shift (data not shown). In figure-9, we showed the formation of 2aa short (254aaG) G-protein due to TCM as compared with 256aaG lineage of HMPV. Similarly, generation of 203aa G-protein due to insertion of one Guanine nucleotide at 6784-position (green circle) creating an early TGA termination codon in OL794386 using upstream ATG initiation codon was suggested (figure-10). Thus, 180nt insertion sequence has duplicated in both Netherland clones but one created 279aa (284aa) G-protein (OL794465) while other (OL794386) created 203aa G-protein. However, a sequencing error was possible as such insertion point was not found in the database (Charlton, 2024)!

**Fig. 7A.**
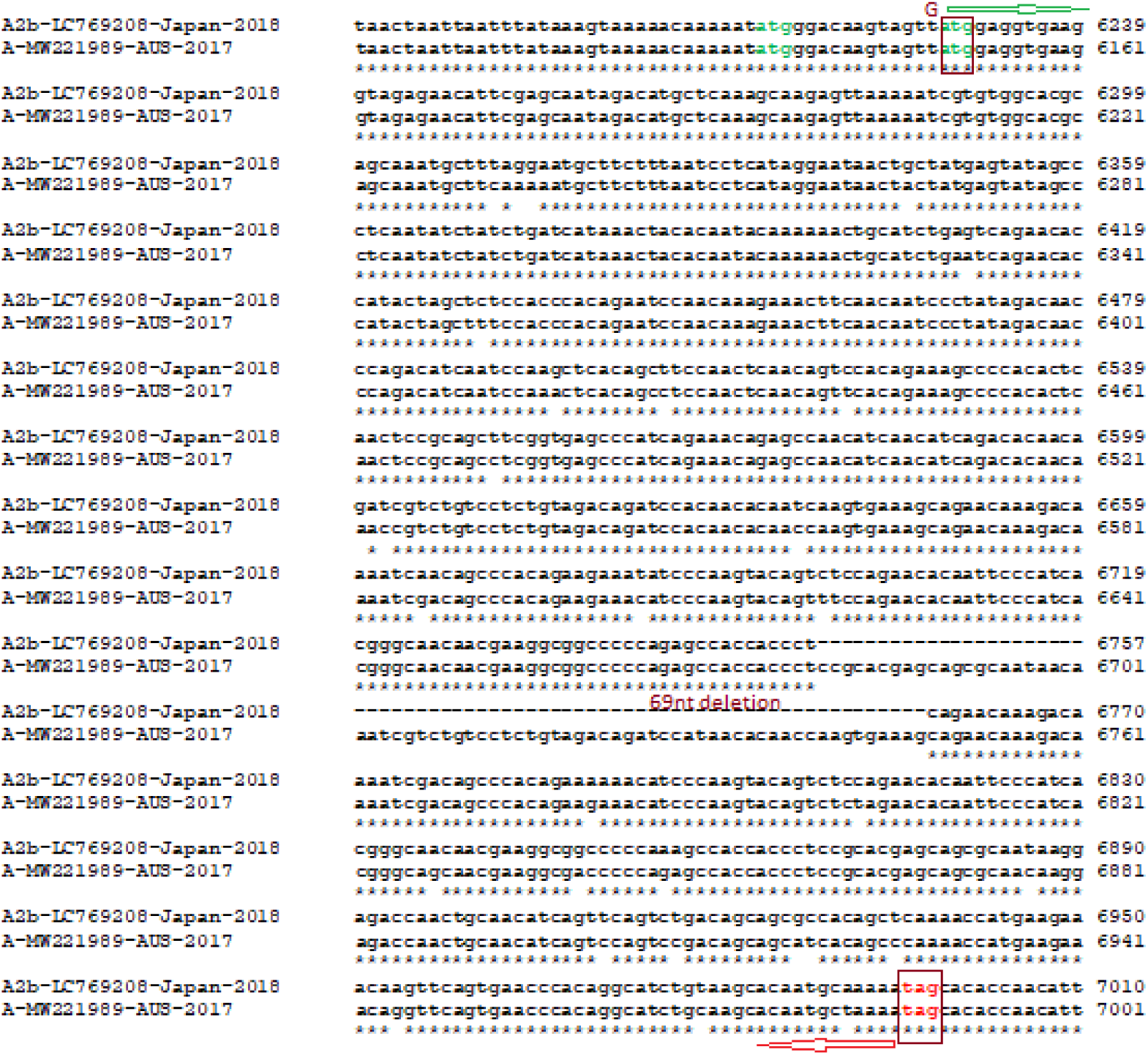
CLUSTAL-Omega similarity between two HMPV sequences (LC769208, Japan and MW221989, Australia) with dissimilarities in G-protein length (279aa vs 256aa) to demonstrate 69nt deletion in the G-gene. Initiation codon (green) and termination codon (red) were boxed. It appeared that such deletion stablished the HMPV as more 256aa G-protein A2-genotype sequences were appeared in the database recently. Thus, duplication of 180nt followed by deletion of 69nt was happened in this Australian clone. The literature however, reported that 180nt and 111nt duplications were independent events and never pointed the deletion event.

**Fig. 7B.**
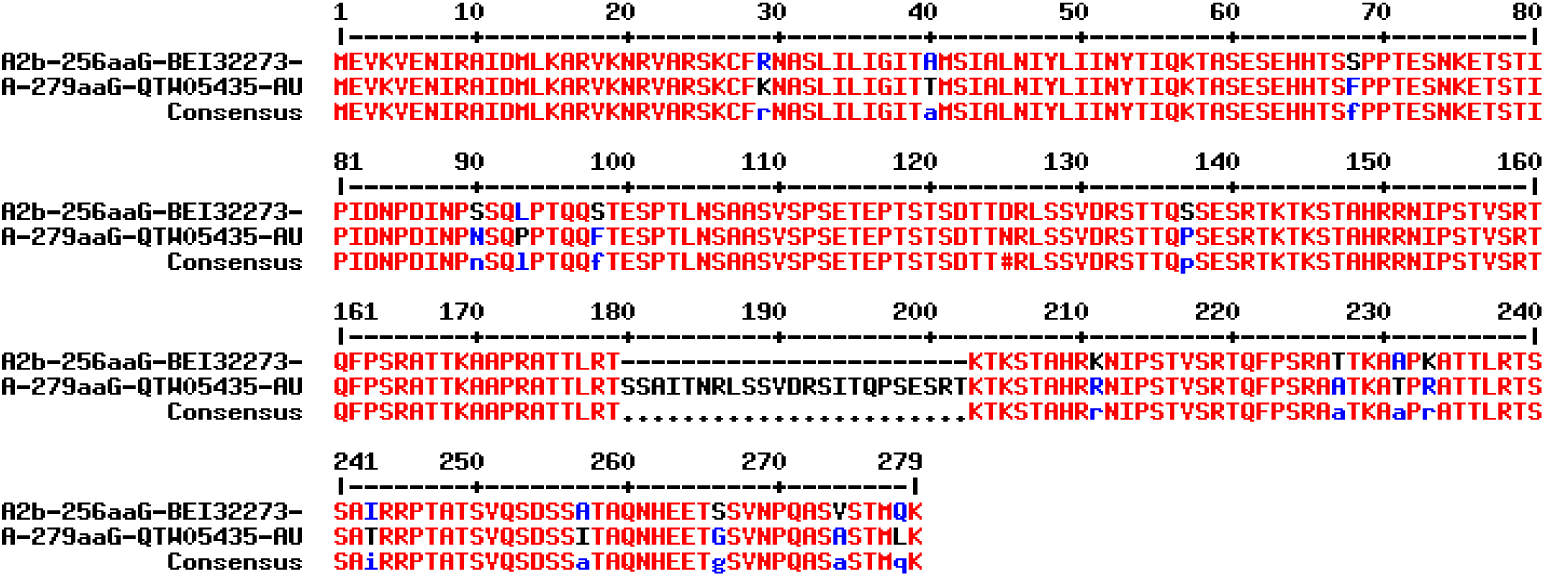
Demonstration of 23aa deletion in the G-protein with 180nt duplication (accession number: LC769208, Japan) compared with complete genome of an Australian clone (accession number: MW221989). The 180-202 amino acids of G-protein (protein id. QTW05435; 279aa) from an Australian clone were deleted as compared to a Japan clone (protein id. BET32273; 256aa).

**Fig. 8A.**
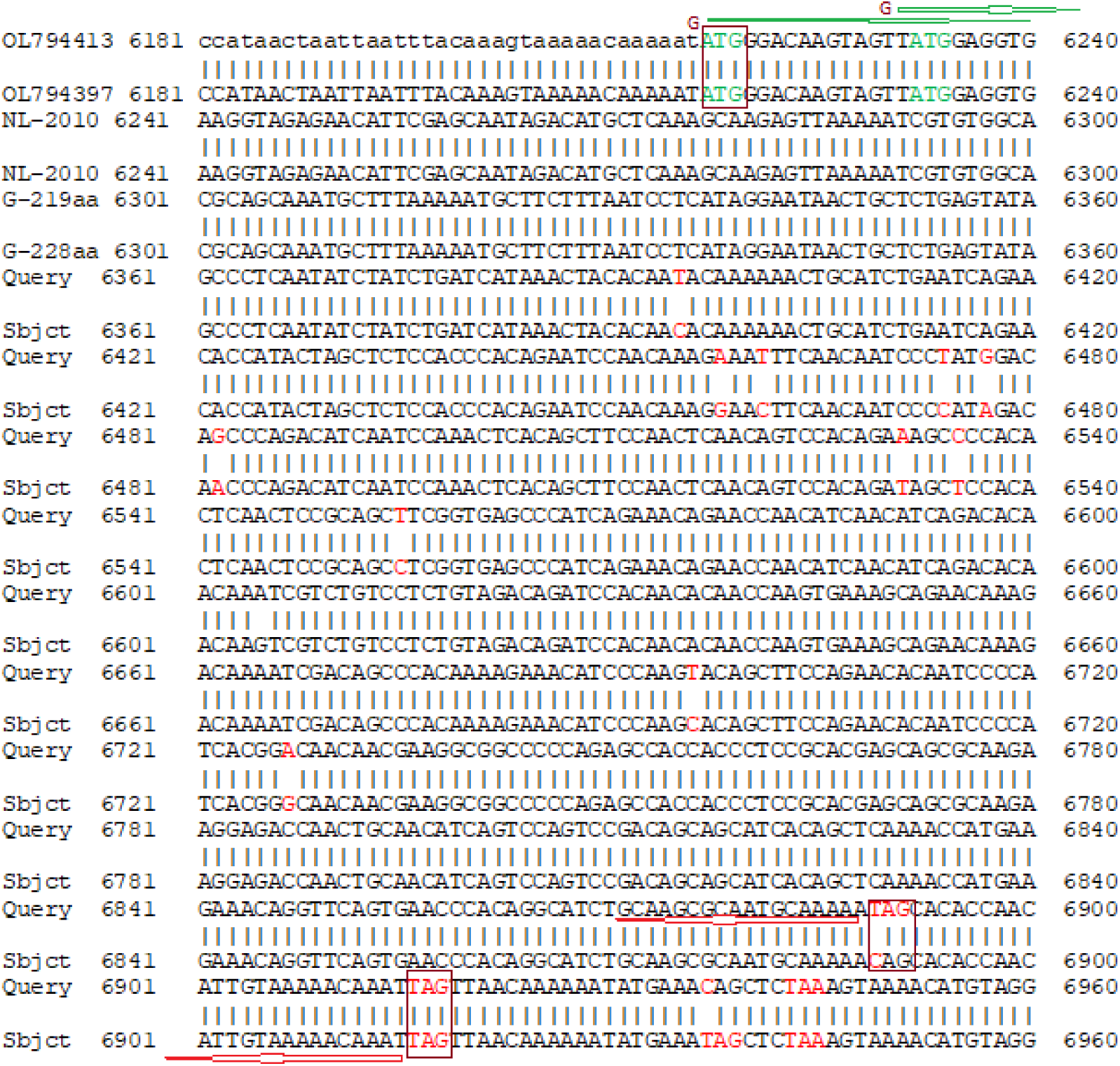
The demonstration of the A-genotype HMPV 219aa (224aa) G-protein into 228aa (233aa) due to point mutation (6889T>C) destroying termination codon (TAG=CAG; OL794397) and extended 228aa G-protein (protein id. WAB07869) was produced from the second TAG termination codon.

**Fig. 8B.**
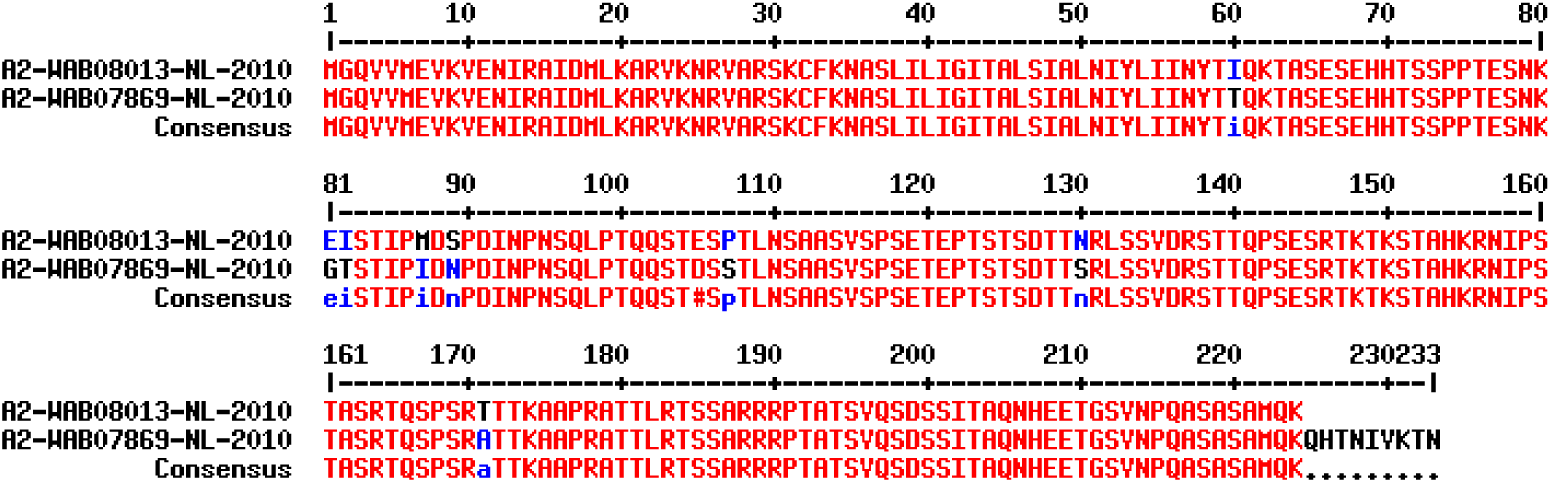
Demonstration of 9aa extended G-protein (233aa, using upstream ATG initiation codon) due to destruction of termination codon as shown in figure-8A. The protein was terminated however from the downstream TAG termination codon. Thus, even both assigned as A2 genotype, there was a difference at the Carboxy terminus and other point mutations. Truly, if we have to assign the HMPV classification according to COVID-19 (SARS-CoV-2), then one has to introduce thousand genotypes as exemplified in A2.2.2 by Devanathan et al. 2025 and Groen et al. 2023.

**Fig. 9A.**
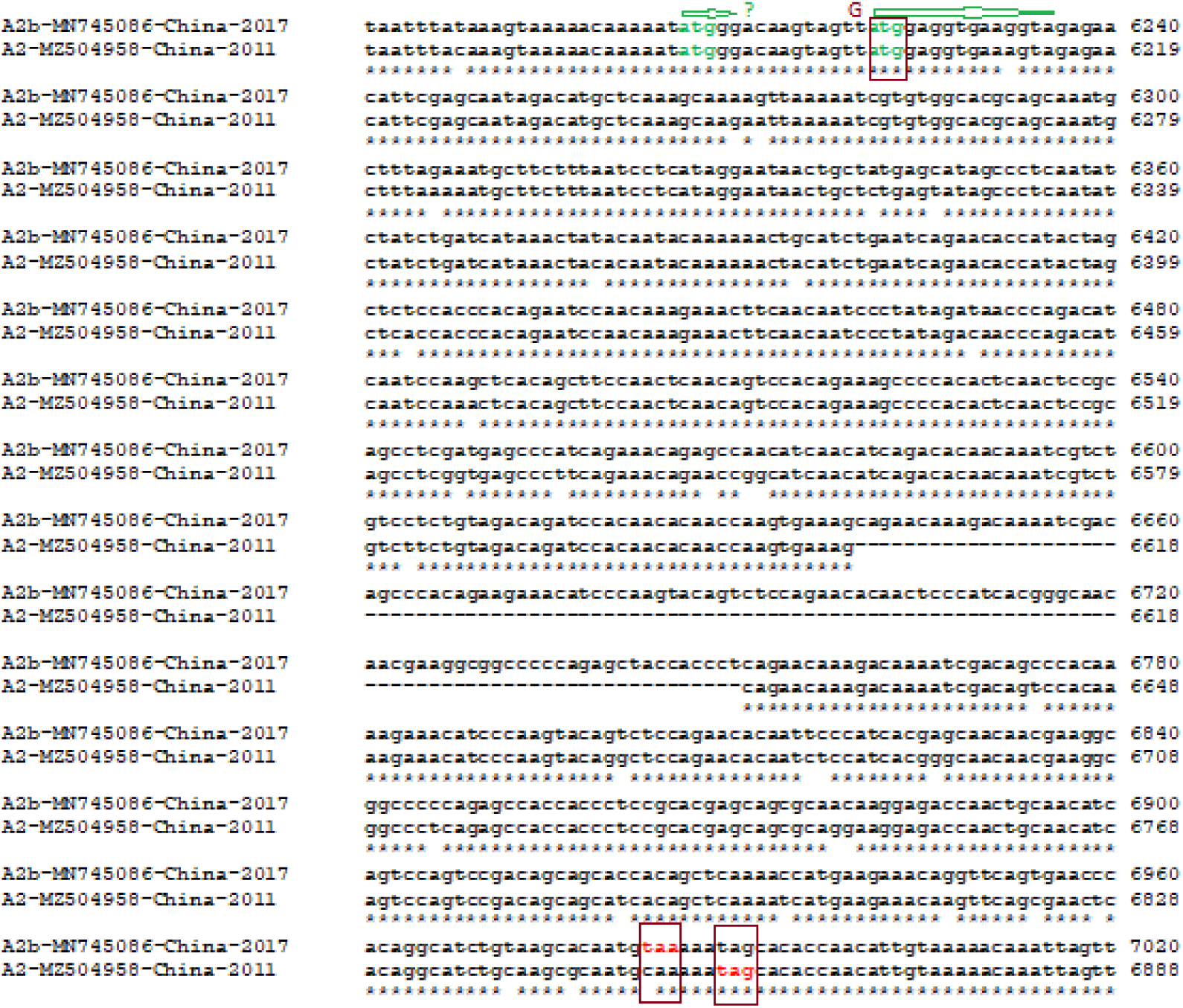
Demonstration of 111nt duplication and C6983T mutation (accession no. MN745086) creating 254aa G-protein creating CAA=TAA early termination in A2-genotype. While China clone (accession no. MZ504958) has no insertion likely early HMPV clone creating 219aa G-protein (A-genotype). The initiation codon (green) and termination codon (red) were boxed.

**Fig. 9B.**
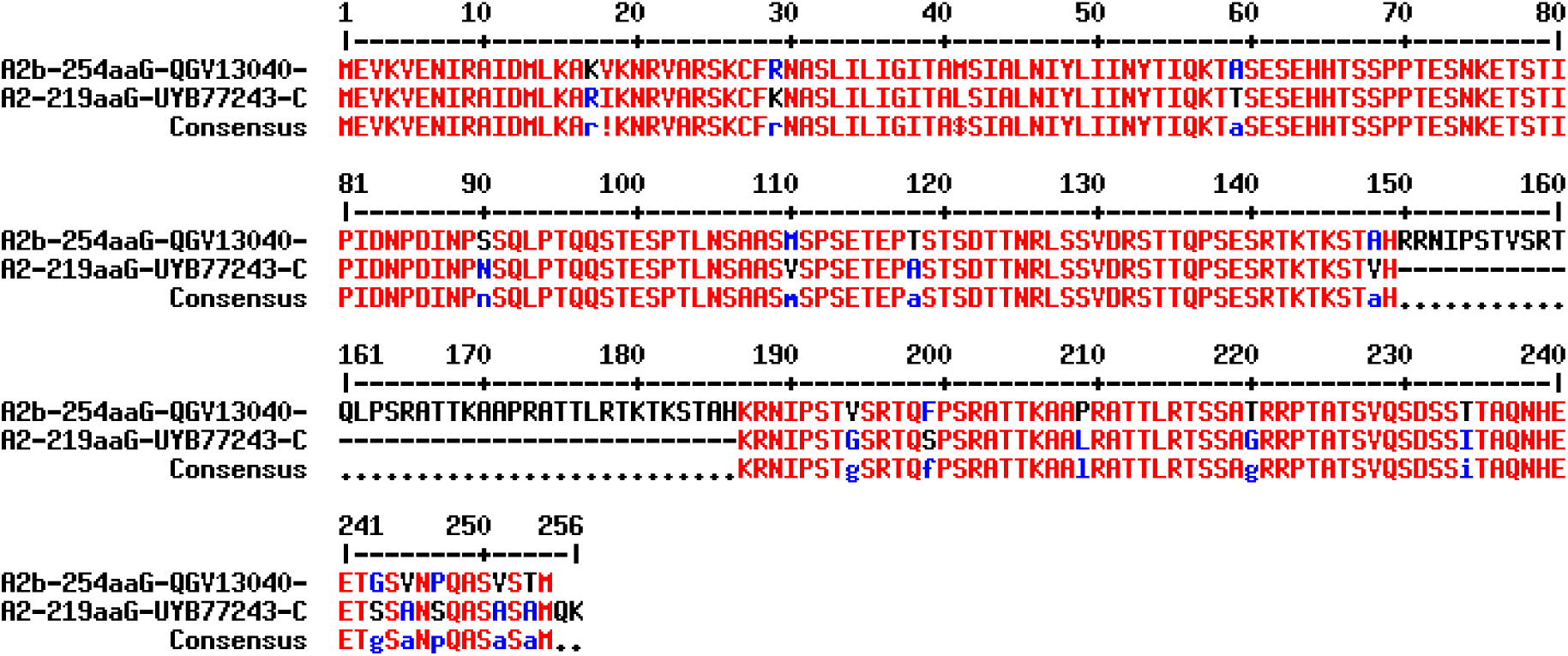
G-protein sequence similarities between two A-genotype (QGV13040 and UYB77243) sequences as described in figure-9A with 37aa insertion and an alternate new termination codon (CAA to TAA) due to C6983T TCM.

**Fig. 10A.**
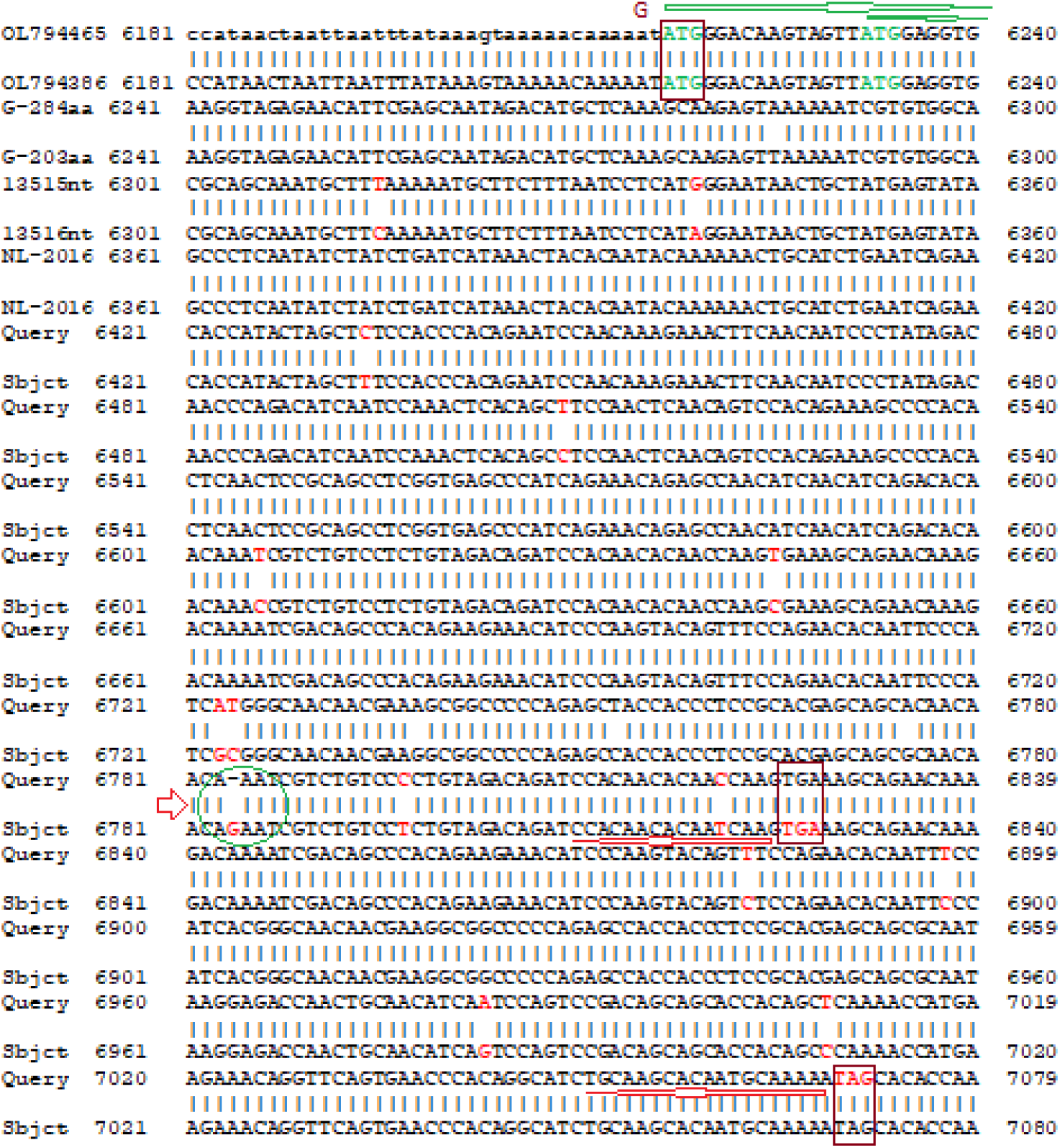
Generation of 203aa G-protein due to insertion of one Guanine nucleotide at 6784 position (green circle) creating an early TGA termination codon in OL794386 using upstream ATG initiation codon. Thus, 280nt insertion sequence has duplicated in both Netherland clones but one created 284aa G-protein (OL794465) while other (OL794386) created 203aa G-protein. However, a sequencing error was possible!

**Fig. 10B.**
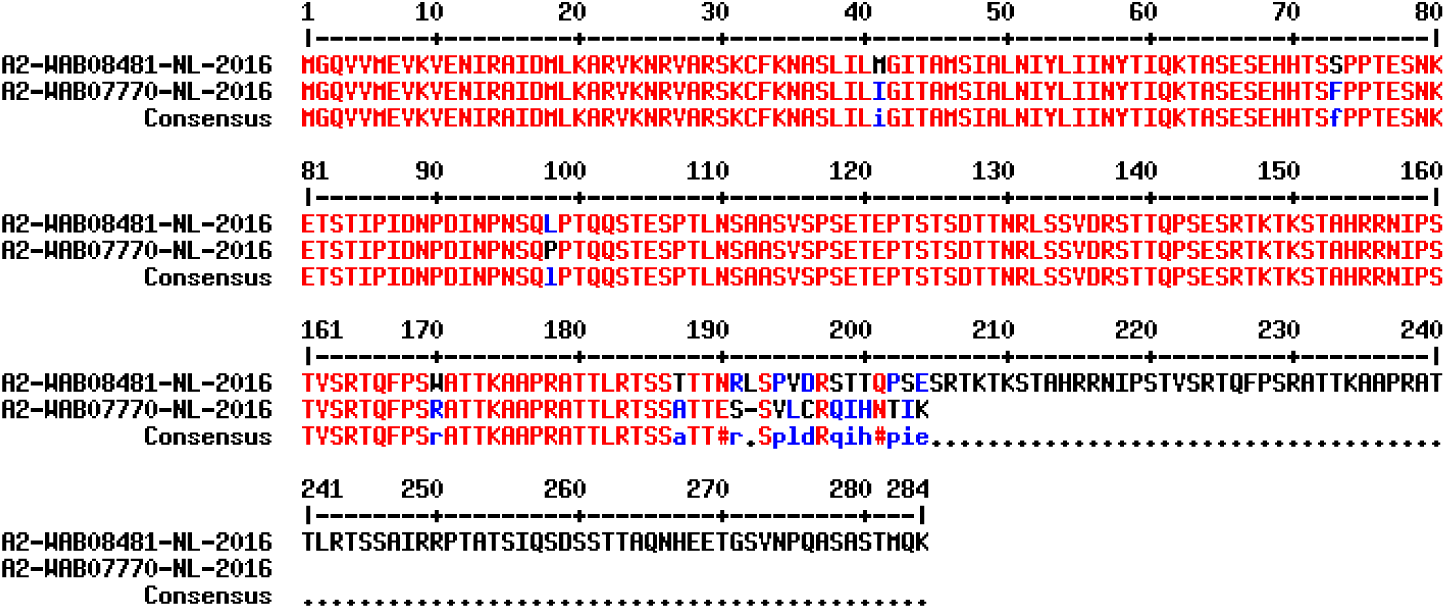
Difference between 284aa (279aa) and 203aa (198aa) two G-proteins from both 280nt duplication HMPV clones from Netherland (OL794465 and OL794386) due to one nucleotide insertion at 6784-position. BLASTP search of carboxy-terminal 23aa (H2N-tlrtssattessvlcrqihntik-CO_2_H) only produced one sequence (protein id. WAB07770) suggesting a sequencing error.

In figure-11, we demonstrated the destruction of termination codon (TCM; TAG=CAG) in G-protein of US A-genotype HMPV genome (PV052163) creating 5aa extended G-protein (261aa) using second alternate termination codon (TAA) as compared with 256aa G-protein A2-genotype clone (PV052139). The generation 237aa G-protein (WAB08643) was formulated in figure-12 locating a TCM in the G-gene but also used ATC extended few amino acids. Here, we aligned A-genotype verses B-genotype explaining less homology. But we yet to determine the genesis point of B-genotype of HMPV! However, we have given an example here favoring 256aaG clone as source of B-genotype lineages and it was explained clearly.

**Fig. 11A.**
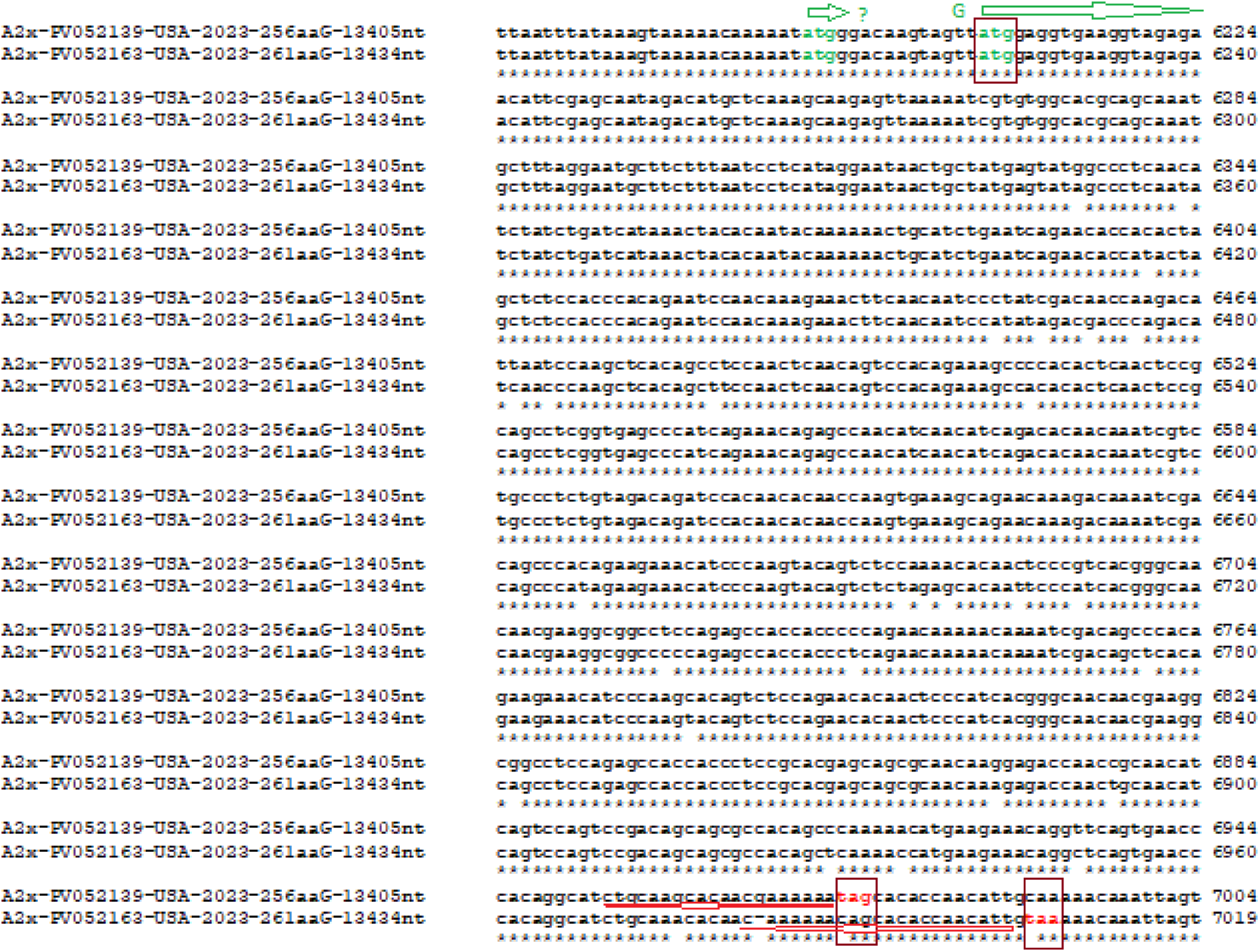
Demonstration of destruction of termination codon mutation (TAG=CAG) in G-protein of US A-genotype HMPV genome (PV052163) creating 5aa extended G-protein (261aa) using second alternate termination codon (TAA) as compared with 256aa G-protein A2-genotype clone (PV052139).

**Fig. 11B.**
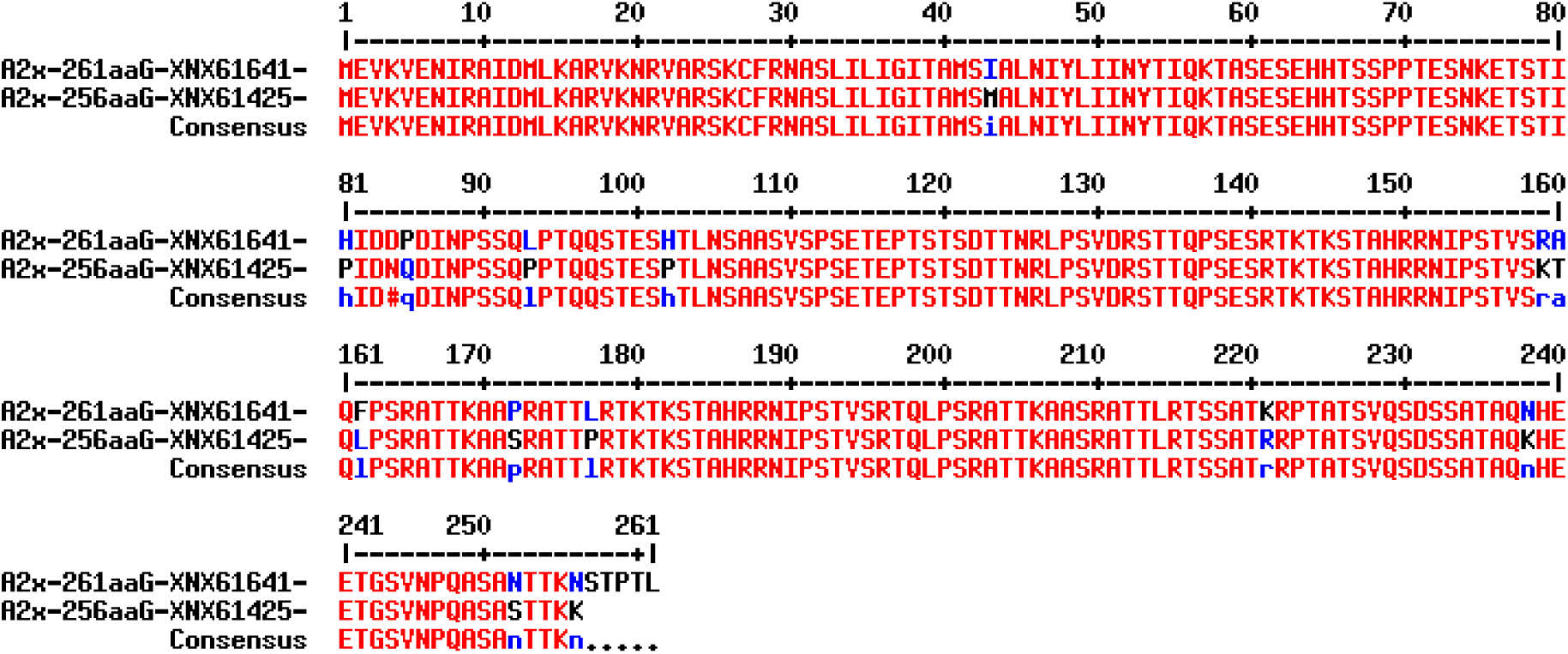
Demonstration of sequence similarity of two G-proteins (261aa vs 256aa) of two US HMPV A2-genotype clones. We suggest the genotypes as A2.256.US.1 (PV052139) and A2.261.US.2 (PV052163).

**Fig. 12A.**
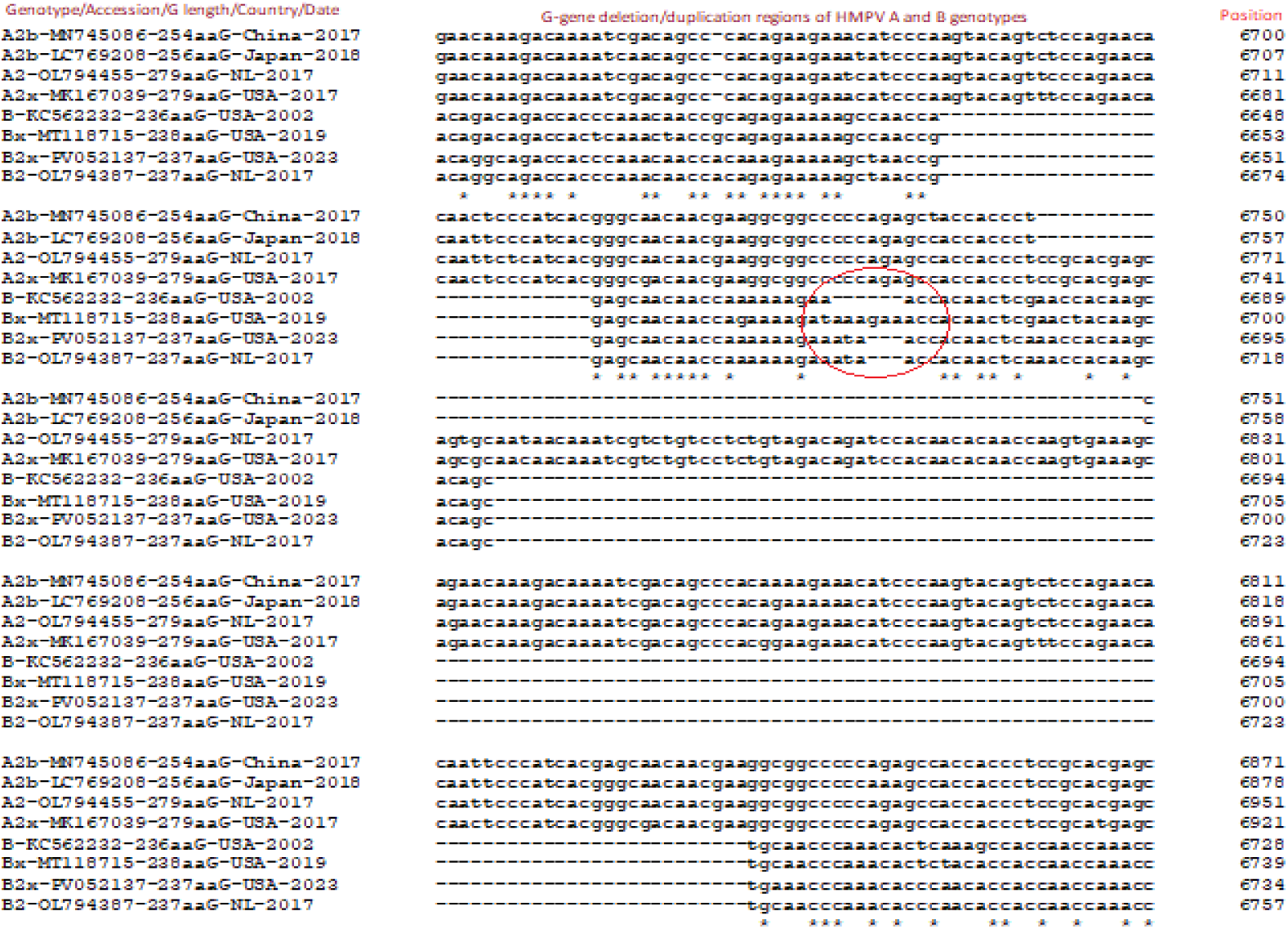
The multi-alignment of genomes of HMPV A-genotype and B-genotype to demonstrate the duplication/deletion G-gene regions of 279aa (OL794455, MK167039), 256aa (LC769208), 254aa (MN745086), 238aa (MT118715), 237aa (PV052137), 236aa (KC562232, OL794445) lengths G-gene regions. The 238aa, 237aa and 236aa clones are B-genotype and minor deletions were located (circled). The AAAGAA deletion in 236aa G-gene removed Lysine and Glutamic acid (KE).

**Fig. 12B.**
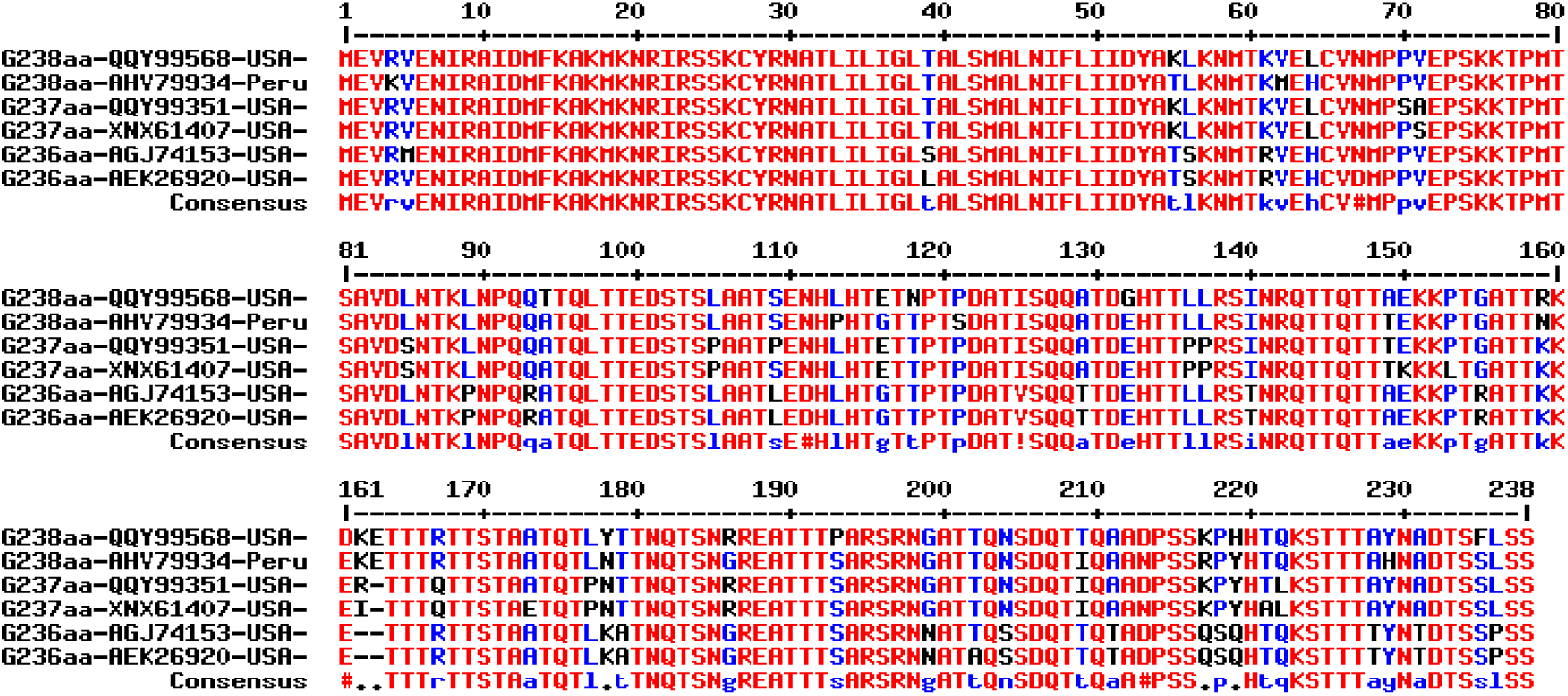
The multi-alignment to demonstrate the one (E) and two (KE) amino acid differences among 237aa and 238aa G-protein HMPV B-genotype clones with respect to 236aa G-protein.

When, we compared the G-gene with same genotype, still we noticed few mutations and demographic changes were also important. When we compared two B1-genotype HMPV sequences obtained from Australia in 2023 and Japan in 2022 respectively. The OY757729 is monopartite sequence and genotype obtained by sequence similarity. Both cases 241aa G-protein were produced. When we BLASTP searched with 5’-gagaactcaa cattttcggc agcaacccca gagggccatc catacacaga gacaattcaa-3’ (6540 position) sequence only two WGS sequences (OY757727, OY757729) with 100% similarity were obtained. But such study with 5’-gagagcccaa cattttcggc agcaacccca gagggccatc catacacaga gacaacccaa-3’ (6540 position) sequence only one WGS sequence (LC769218) was obtained. This suggested that minor variation among the same genotypes always happened in the G-gene of HMPV (data not shown). However, sequence error possible here (Goudey et al. 2022). The closest WGS sequences were OP904127 (Australia-2018, B-genotype) and PQ634885 (USA-2021, Bx-genotype).

In figure-13A, we showed another example of TCM where T6990C mutation (TAG=CAG) generated 265aaG extended using ATC downstream (LC275891=265aaG and LC67155=256aaG) but few WGS sequences were found only in the database. The C-terminal changed from VSTMQK (PP947648, OP904022) to MAKTN (LC275891, PP947641). However, major 256aaG ends with VSTMQN (OP904007, OP904030, LC671555, LC769209) or ASTMQK (PP947557, PP947617, PP947676). The A2.256.1 and A2.265.1 may be the genotypes here but country name must be used. As for example, G-protein of HMPV of recent US clones had preferably ends with ASTMQK and thus, here we should write A2.256.USA.1. If you got 100 sequences from a study, you could be used A2.256.USA.1 to A2.256.USA.100. When you multi-aligned the 100 sequences, then you would be able to further classify sub sub-genotypes like A2.256.USA.1.1 (PP947557) or A2.256.USA.1.2 (PP947612). If you want to mention the individual mutations, that is also possible. Even such nomenclature may be matched in different workers in the same country without any harm if a consensus was achieved in international symposium.

**Fig. 13A.**
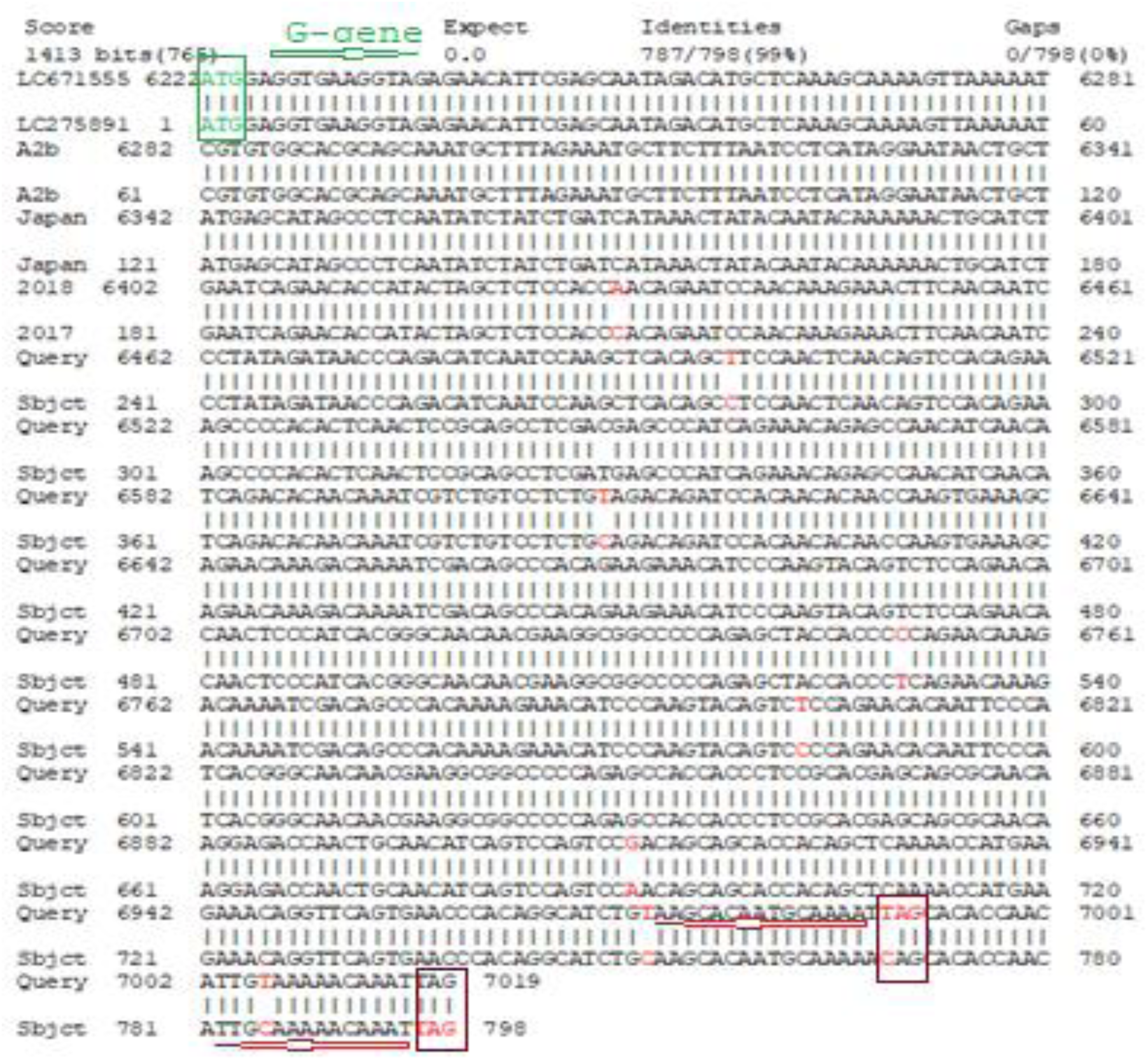
Demonstration of T6990C TCM to generate 265aaG using ATC downstream (LC275891=265aaG and LC67155=256aaG) but few WGS sequences were found only in the database.

**Fig. 13B.**
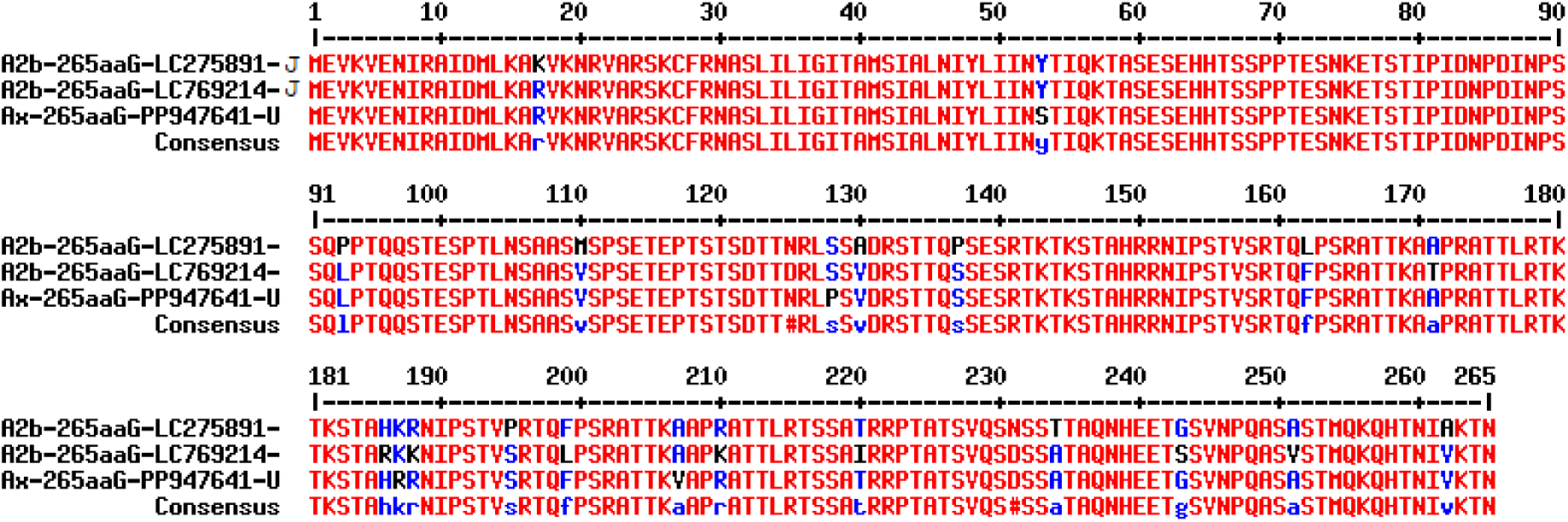
Differences in 265aa G-protein of A2-genotype due to TCM. We found only three such WGS clones in the database and had few mutations among themselves. The 9aa extended G-protein here ended with NI(V/A)KIN instead VSTMQN or ASTMQN as found in most 256aaG A2-genotype clones.

Again, we dared to compare using BLAST2 alignment between 236aaG A-genotype (KC403976) vs 236aaG B-genotype (KC562228) (figure-14). Both used similar initiation and termination codons but from what ancestral both were generated? The 52% sequence similarity was observed between two G-genes. The WGS similarity was found 84.1% between A-genotype and B-genotype HMPV suggesting strong heterogeneity in G-gene region.

**Fig. 14A.**
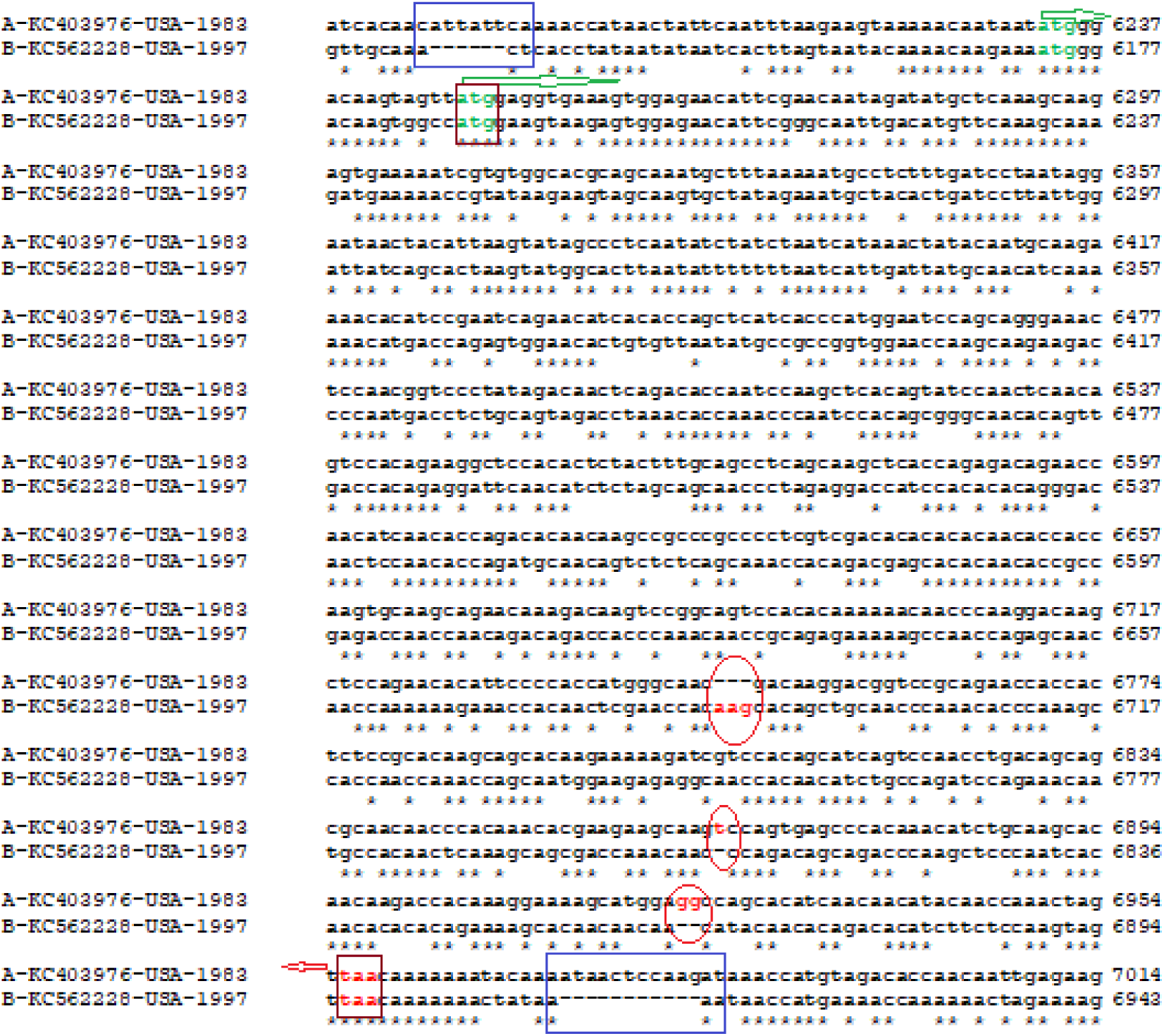
BLAST2 alignment between 236aaG A-genotype (KC403976) vs 236aaG B-genotype (KC562228). Both used similar initiation and termination codons but from what ancestral both were generated? The 52% sequence similarity was observed between two G-genes. The WGS similarity was found 84.1% between A-genotype and B-genotype HMPV suggesting strong heterogeneity in G-gene region.

**Fig. 14B.**
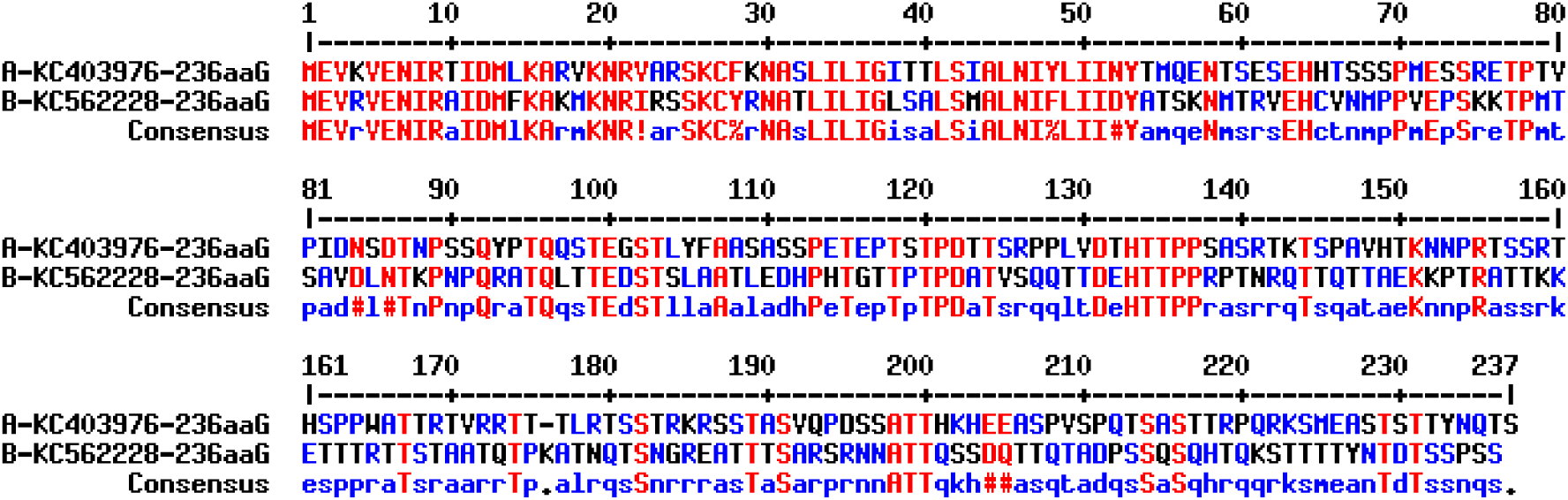
Huge differences between 236aaG protein of A-genotype verses B-genotype. Does 236aaG HMPV clones ancestral HMPV clones? Only 35% sequence similarity between two G-proteins of A-genotype and B-genotype.

In figure-15, we demonstrated the generation of 237aaG (242aaG) in B-genotype of HMPV. The HMPV genomes with 279aa (OL794455 and MK167039), 256aa (LC769208), 254aa (MN745086) and 237aa (PV052137 and OL794387) G genes were aligned to demonstrate 153nt deletion in genomes with 237aa G-protein. However, if such lineages produced from 256aa or 279aa G-protein lineages was not clear. It is an endeavor to get clue to idea of genesis of B-genotype of HMPV. The accepted values of duplication lengths were always found 180nt and 111nt as described by many workers (Saikusa et al. 2018; Parida et al. 2023). However, termination codon was different between A-genotype and B-genotype. An early termination codon (CAA=TAA) was generated due to C6870T mutation in 254aa lineage (MN745086) causing two amino acids short G-protein as compared to 256aa G-protein lineage. The WAB0779-Netherland and XNX61407-USA 237aaG had eight mutations due to demographic differences (figure-15). we had also checked the difference in G-gene of A1 genotype (protein id. AEK26893, 236aa, USA) and A2 genotype (protein id. AGJ74188, 236aa, Australia) of human metapneumovirus. Thus, both sequences (JN184399=13389nt and KC562236=13340nt) had no duplication in the G-gene and were given new genotyping as A.236.1.USA genotype and A.236.2.AUS respectively (data not shown).

**Fig. 15.**
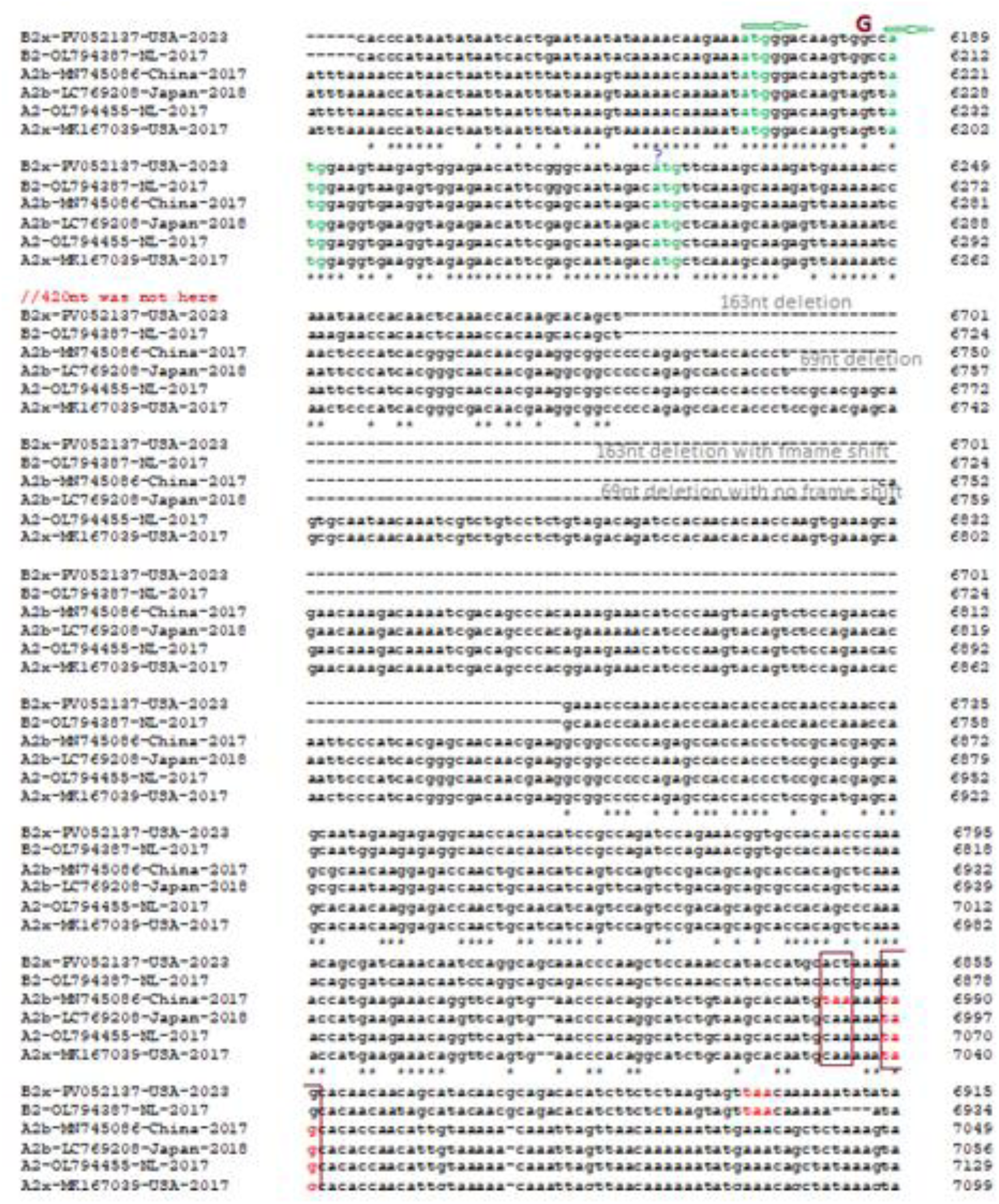
Generation of 237aa (242aa) B-genotype of HMPV. The HMPV genomes with 279aa (OL794455 and MK167039), 256aa (LC769208), 254aa (MN745086) and 237aa (PV052137 and OL794387) G-proteins were aligned to demonstrate 153nt deletion in genomes with 237aa G-protein. Thus, if such lineages produced from 256aa or 279aa G-protein lineages was not clear. However, termination codon was different between A-genotype and B-genotype. An early termination codon (CAA=TAA) was generated due to C6870T mutation in 254aa lineage (MN745086) causing two amino acids short G-protein as compared to 256aa G-protein lineage. The WAB0779-Netherland and XNX61407-USA 237aa HMPV G-proteins had eight mutations due to demographic differences.

Again, we checked the difference in nucleotides or amino acids between related G-gene regions of HMPV genomes (MZ851993 and OP904134; A-genotype) one from Australia and the other from China. Both cases 256aa of G-protein was produced and Australian clone should be A2.256.AUS.1 genotype while China clone should be A2.256.CN.2 genotype (data not shown). The Australian scientist claimed as A-genotype or B-genotype only without doing any further sub-classification. Whereas Netherland scientists preferred as A1/A2 and B1/B2 classification. However, since 2015 most US scientists did not mention any genotype during their GenBank submission and similar notion was lately followed by Chinese scientists. The Japanese scientists followed A2a/b/c and B1/B2 classification as followed by Indian scientists. However, I did not find any HMPV WGS in the GenBank database from India!

Similarly, we checked two same A2b genotypes were genetically different and must be renamed genotyping! The Japan lineage (LC671558) was found 111nt deletion in G-gene and China lineage (OP358287) had 111nt duplication. We have given the new genotype name as A.219.1.JP.1 genotype to Japan lineage while A2.256.1.CN.1 genotype to China lineage. The termination codon TAG remains same producing one G-protein with 256aa (protein id. WGF20462) and other with 219aa (protein id. BDE17504) (data not shown).

The sequences of two genotypes HMPV had huge variation. So, why we did not write the genotype during GenBank submission? As for example, we also checked the BLAST2 alignment between 236aaG A-genotype (KC403976; A.236.1.USA.1) vs 236aaG B-genotype (KC562228; B.236.1.2.USA.1) (data not shown). The initiation and termination codons were found same. The 52% sequence similarity was observed between two G-genes. The WGS similarity was found 84.1% between A-genotype and B-genotype HMPV suggesting strong heterogeneity in G-gene region. Truly, SH-gene located upstream was also heterogeneous. The F-gene and L-gene were very conserved and also the N-gene and P-gene to some extent. A mere 35% sequence similarity between two G-proteins of A-genotype and B-genotype (data not shown). We faced problem during alignment among A-genotype and B-genotype sequences due to higher mismatch. Fragmentation occurred using BLAST2. However, CLUSTAL-Omega produced quite expected data. We found higher differences of G-gene of HMPV from Australian patients between A-genotype (OP904134) and B-genotype (OP904034) (data not shown). CLUSTAL-Omega alignment gave data but 111nt deletion was not there instead maximum 102nt deletion in three phases were demonstrated likely due to higher percentage of mutations and termination codons were different between A-genotype and B-genotype. The G-protein of A-genotype is 256aa while B-genotype is 237aa. However, deletion region predicted at the carboxy-terminus but hard to conclude (data not shown).

Thus, the RNA sequence as well as amino acid heterogeneities of G-gene between A and B genotypes were maximum and TCM produced the varying lengths of attachment proteins. Now, we will consider more carefully the amino acid variation between similar genotypes of HMPV. As for example, huge variation of amino acids in G between A2 (G=AGJ74188, 236aa, 2004) and A2b (G=WGF20462, 256aa, 2019) of HMPV. Thus, there genotype must be different. The A2 to A.236.AUS.1 genotype (KC562236) and A2b to A2.256.CN.1 (OP358287) genotype was proposed here (data not shown). We also checked the difference between B2 genotype 237aaG clones one from Netherlands (OL794483, B2, 2017) and other from Australia (OP904034, B, 2019). The mutational differences could be managed by writing B2.237.AUS.1 whereas we can write more correctly as B2.237.AUS.1:I108N:P121L:L135P. The WAB08643 used upstream ATG initiation codon for G-gene and designated as 242aaG (data not shown). Similarly, we disclosed a strong variation between two A2b genotype G-proteins (BDE17504, LC671558, 219aa, Japan-2018 and WGF20462, OP358287, 256aa, China-2019) and such strong variation at the carboxy-terminus must be justified by new genotyping as A.219.1.JP.1 and A2.256.1.CN.1 (data not shown). Further, we found amino acid sequence variation between two A-genotypes 236aaG proteins from USA, one being A1 and other described A2. But our genotyping nomenclature A.236.1.US.1 and A.236.2.US.1 may be optimistic and you can put the mutations also (data not shown). Important of this article perhaps reside if International HMPV scientists gather in a conference and disclose a consensus method for WGS of all HMPV sequences. Similar way, we noticed the difference between an Australian (KC562236, 13340nt) and Argentinian (DQ362949, 711nt) 236aa G-proteins of HMPV clones showing demographic differences (Galiano et al. 2006).

The recently, we found huge database submission of 256aaG lineages from USA. The terminal few amino acids of multi-alignment of 256aa G-proteins from different A-genotype HMPV clones was demonstrated in figure-16A. We found that the ASTMQK and VSTMQK were maximum ends of G-proteins with 256aa. While in figure-16B the phylogram disclosed a strong demographic relation among 256aa G-proteins with different mutations. We did not find any good relation with mutations in the 256aaG with country or time.

**Fig. 16A.**
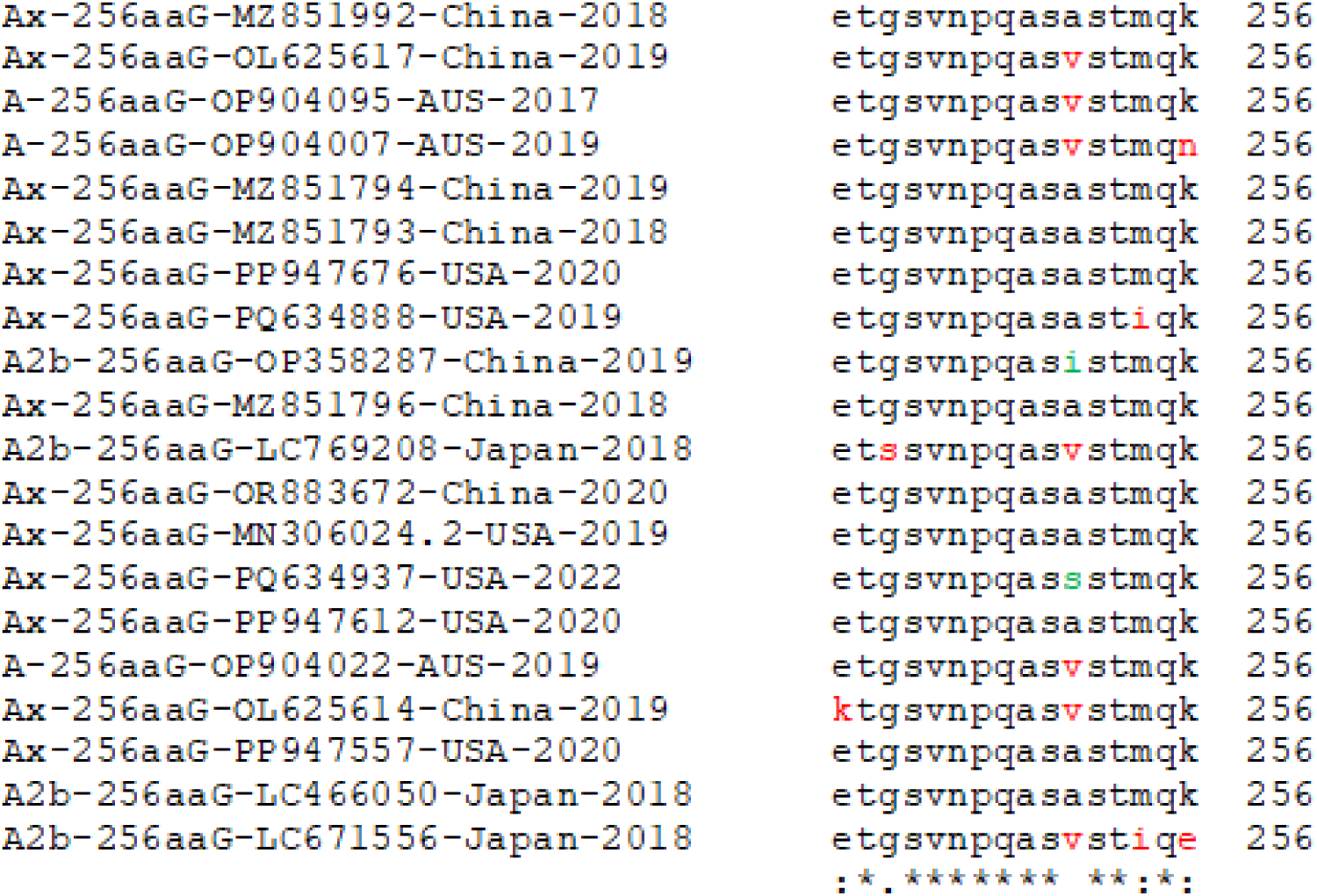
The multi-alignment of 256aa G-protein from different A-genotype HMPV clones. Only C-terminal few amino acids were shown here. ASTMQK and VSTMQK were maximum ends of G-protein with 256aa.

**Fig. 16B.**
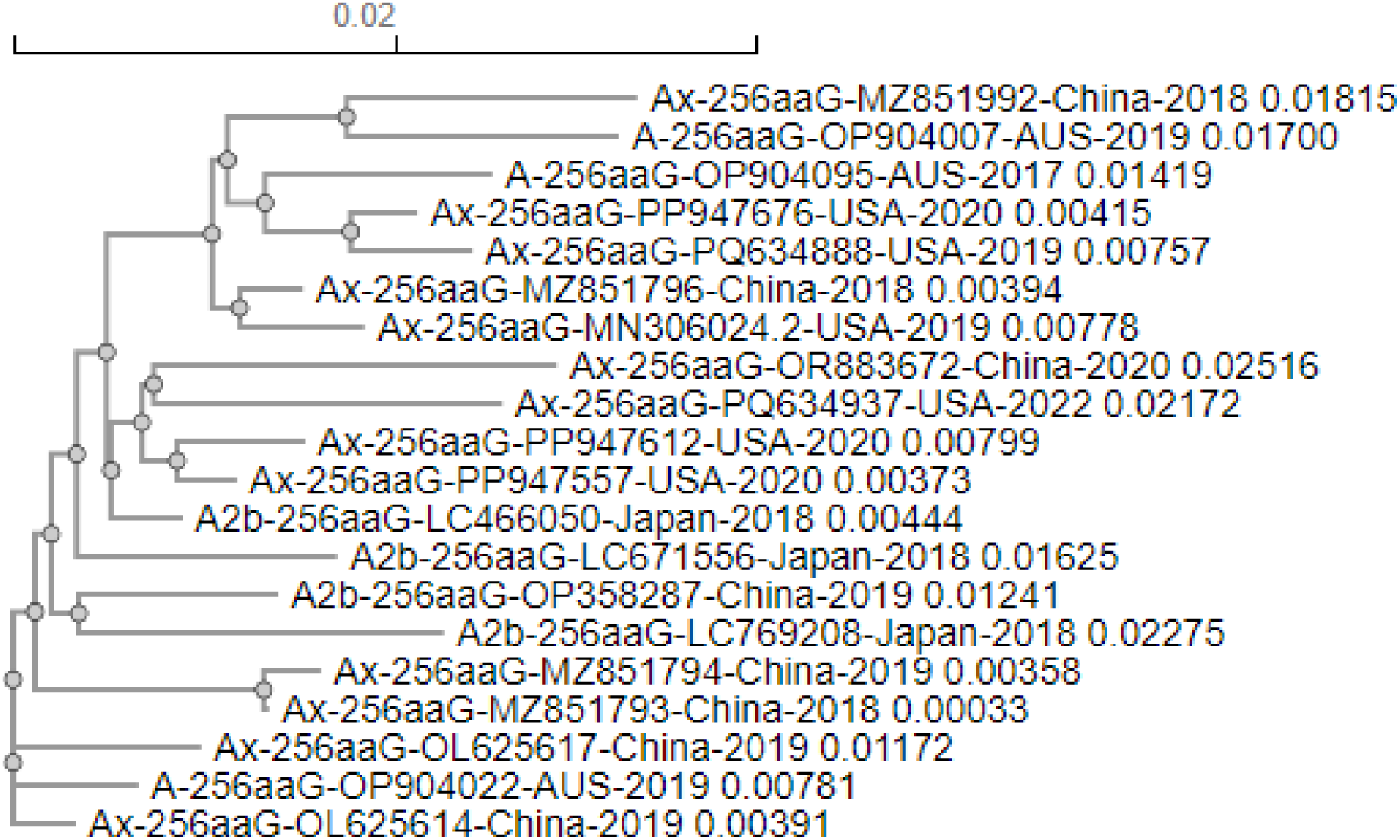
The multi-alignment and phylogeny to check the demographic relation among 256aa G-proteins. We did not find any good relation with mutations in the 256aaG and country or time.

In figure-17A, we described the heterogeneity among four 219aaG proteins isolated from Netherland patients with different time. The data indicated that no 100% similarity, all four lineages were assigned A2-genotypes. Thus, our genotyping nomenclature like A.219.1.NL.1, A.219.1.NL.2, A.219.2.NL.1 and A.219.3.NL.1 are surely good. Similar way we disclosed a phylogram of 219aaG proteins from different countries with different genotypes (figure-17B). The different clusters indicated their name must be different. As the assigned A, A1 and A2 genotypes were meaningless here for similar 219aa G-gene, we controlled the method of genotyping by A-genotype with sub-classification further and denoted by country and amino acid number like A.219.1.NL.1.

**Fig. 17A.**
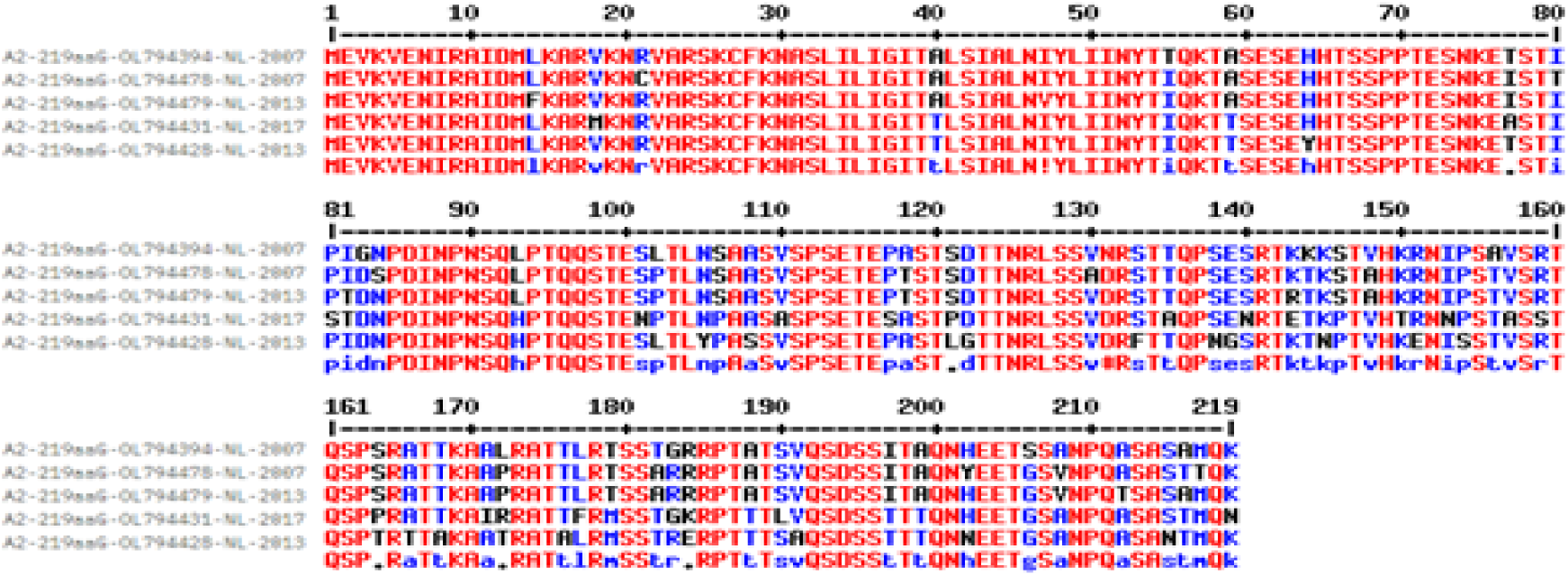
The heterogeneity among four 219aaG proteins isolated from Netherland patients with different time. The data indicated that even no 100% similarity, all four clines were assigned A2-genotypes. Thus, our nomenclature like A.219.1.NL.1, A.219.1.NL.2, A.219.2.NL.2.1 and A.219.3.NL.1 are surely good.

**Fig. 17B.**
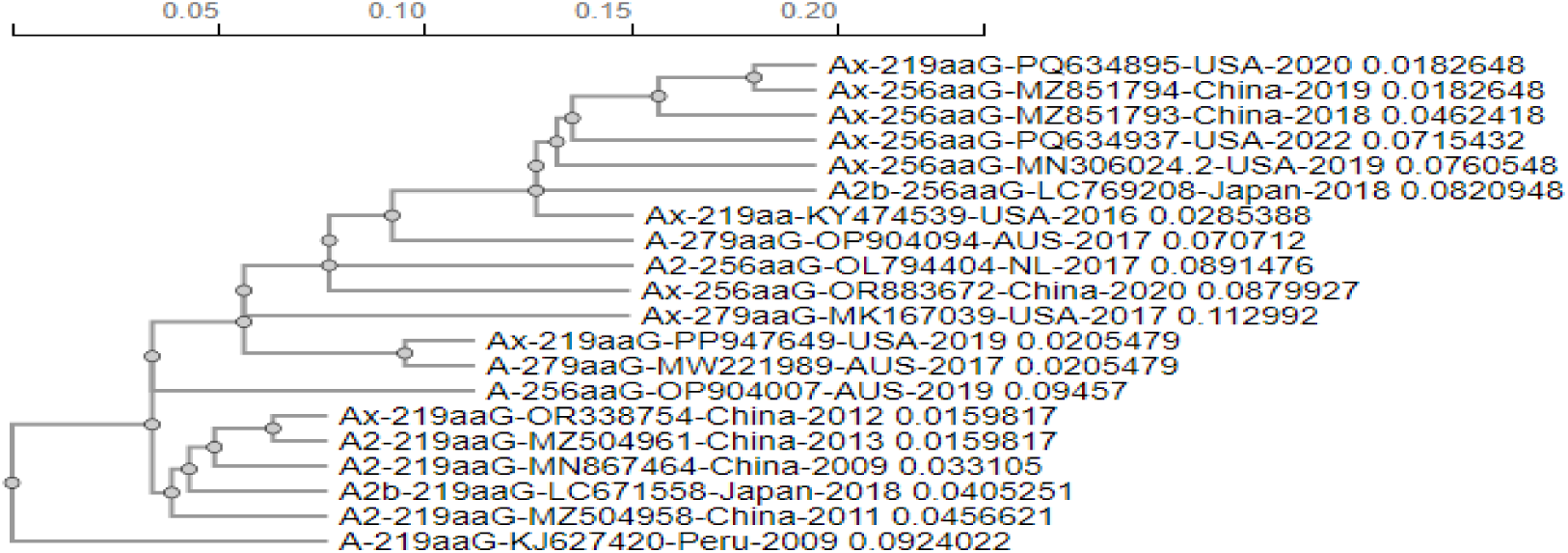
Phylogram of 219aaG protein from different countries with different genotypes. The different clusters indicated their name must be different. As the assigned A, A1 and A2 genotypes were meaningless here, we controlled the method of genotyping by A-genotype with subclassification further and denoted by country and amino acid number like A.219.1.NL.1.

As I repeatedly disclosed that no full length HMPV genome from India (accession numbers: MN410-MN410907 and OR135588-OR135607). But what would be the status of genotyping in India? We checked the data characterizing of A2c genotypes partial G-proteins from India. Data indicated that N90S mutation was the best identity for Indian 256aaG strains of HMPV as compared to partial 279aaG lineages. However, another group from India disclosed the major genotype as A2b while studying F-gene of HMPV (Choudhury ML et al. 2014). However, the F-protein Indian A2b-genotype strains as compared to other A, A2 and A2b genotypes as reported worldwide had no genotype based specific mutation being a conserved protein (data not shown).

We wonder if any variation in the promoter of G-gene. We checked using extensive multi-alignment of many HMPV complete genomes of varying genotypes that indeed there was a bias of differential promoter using between A and B genotypes. We found AATTAATT, ATTTAATT, and AATAATAT for A-genotype and TAATATTAAT for B-genotype promoters (data not shown). However, unless you perform functional assay classical TATATAAT box may not be a functional promoter for RdRP. In N-gene of COVID-19, we reported a similar variation in the promoter among different lineages of coronaviruses (Chakraborty, 2023).

In figure-18, we pointed our method of genotyping so that people can understand the simplicity of such genotyping of HMPV. We happy to disclose another simple method to differ A-genotype of HMPV with B-genotype of HMPV. We found that 3’-UTR 20nt oligonucleotides 5’-TGATAAAATAGATGGCAACT-3’ sequence was not located in the B-genotype of HMPV. Thus, a simple blast search of your WGS with this oligonucleotide will correctly predict whether A-genotype or B-genotype. In truth, we first did 60nt BLASTN search resulting all sequences from A-genotype. When we used only 20nt sequence, many higher eukaryotic sequences were obtained with 100% similarity. As for example, *Monacha cantiana* (snail; ch-14), *Aplidium turbinatum* (sea squirts; ch-1*), Haliotis tuberculata* (sea snail; ch-12) and *Pempelia palumbella* (moth; ch-18) were highlighted with HMPV A-genotype sequences (data not shown). Another important observation was the B-genotype SH-protein had a single amino acid deletion (F58) that was not located in the SH-protein of A-genotype.

**Fig. 18.**
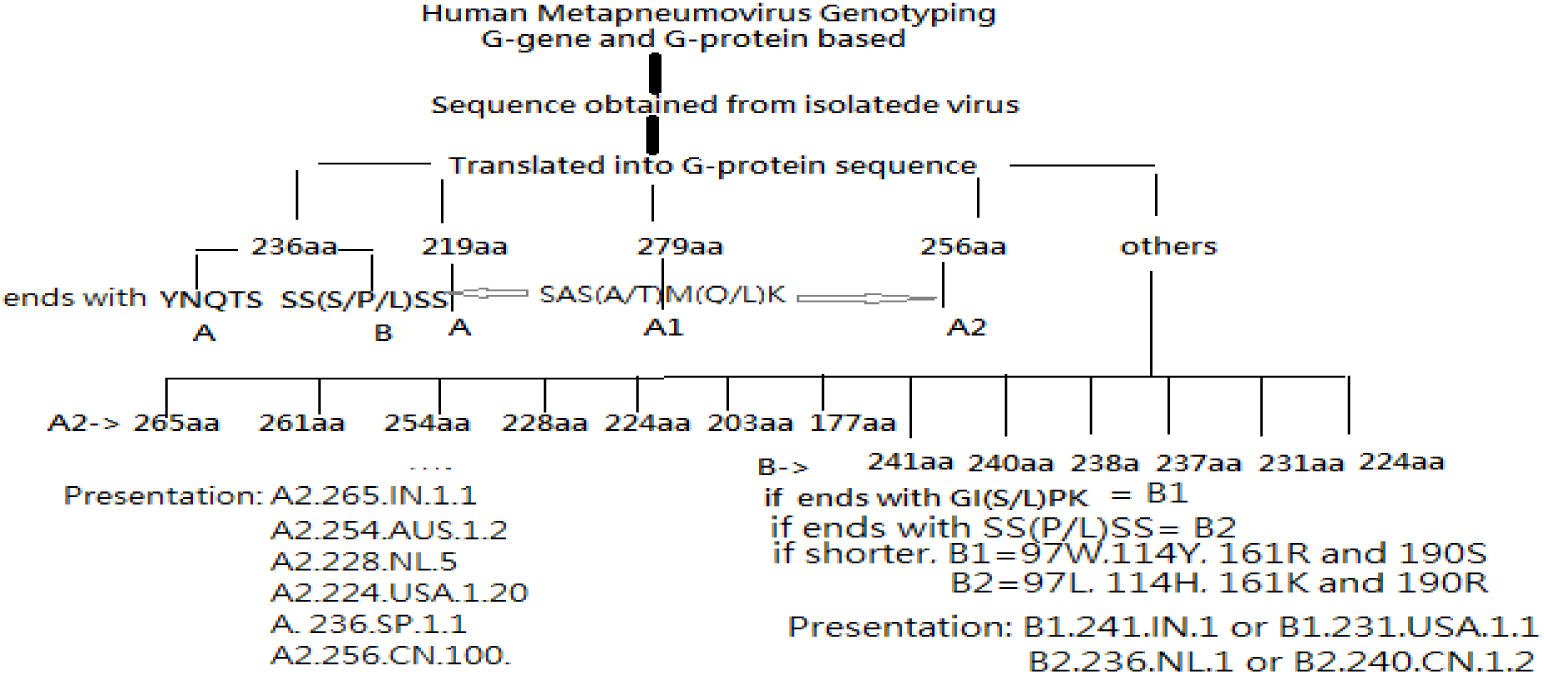
Flow diagram to know HMPV genotype if G-protein sequence was known. IN=India, NL=Netherland, CN=China, USA=Unites States of America, SP=Spain, JP=Japan and so on. The country name was used to minimize the G-protein variation like we had seen African States.

## Discussion

The outbreaks of two genetic lineages (A1/A2 and B1/B2) of HMPV as well as genetic sub-lineages (A2a, A2b, A2c or A2.2.2) occurred worldwide whereas recent China outbreaks might be significant. In 2018, HMPV-associated hospital admissions among children amounted to 643,000 with 16,100 (hospital and community) HMPV-associated ARTI deaths. In Australia, the incidence of HMPV infection surged three-fold in 2021 compared to the period of 2017∼2019 (Ji et al. 2024; Chow et al. 2016). Nevertheless, the epidemiological and clinical characteristics of HMPV infections have shown disparities following the COVID-19 pandemic. The RT-PCR testing was utilized to detect HMPV infection in India but due to poor sequencing standard the scenario of HPMV and HRSV infections was never critically studied in India (Parida et al. 2023). The important error we have found that a Peru HMPV B-genotype clone with designated as A-genotype in accession number KJ627432. Similar mistake was found in USA A1-genotype clone demonstrated in the database as B1-genotype in accession number PP086008. The multi-alignment of G-proteins from partial sequences was critically assessed but failed to draw a conclusion being incomplete sequences.

The HMPV dominant lineages were changed from year to year in different countries and G-gene complexities were seen for genotyping (Biacchesi et al. 2003; Kim et al. 2016; Tulloch et al. 2021). Recently Devanathan et al reported an outbreak in South India from November 2022 to March 2023, with 56 of 129 cases (43.1%) positive for hMPV, making it the most prevalent pathogen, followed by H3N2 influenza outbreaks (n = 34, 26.2%). Sadly, no Indian WGS HMPV sequence was found in the GenBank Database! The panic of COVID-19 is vivid. I read the newspaper for HMPV outbreaks in China followed by India. I decided to study the virus through bioinformatics approach. The G-gene and G-protein multi-alignment data were horrible as no interpretable alignment and no proper phylogenetic clue if incomplete genes were used for analysis. The A, A1 and A2 aligned together while B1 and B2 genotypes were aligned quite satisfactory. So, I first tried to group the G-proteins on the basis of amino acid lengths and also used complete sequences only, where I got a clue to optimise the data with correct interpretation (figure-4). The 236aaG lineages appeared early, then 219aaG and then 279aaG (180nt duplication) followed by 256aaG (69nt deletion or 111nt duplication) being all A-genotypes. While 236aaG B-genotype appeared first and then 242aaG, 240aaG, 237aaG and so on. The smaller version of G-proteins (217aa, 224aa, 228aa) are always there due to TCM. However, extended G-proteins like 261aaG and 265aaG could be seen due to extension using downstream second termination codon (figure-11 and figure-13). The Netherland scientists deposited data using second upstream initiation codon extending five amino acids (279=284aa; 256=261aa; 237=242; 236=241aa and so on). Such G-protein sequences were adjusted removing five amino acids from N-terminus during multi-alignment. Most remarkable finding was that the TCM similar to SARS-CoV-2 XBB.1 variant as well as XBB.1.5.1-XBB.1.5.100 subvariants where ORF8 protein 8^th^ codon GGA=TGA mutation caused no ORF8 protein in such variant of COVID-19 lowering immunogenicity (Chakraborty, 2023). We showed in figure-2A/B the duplication of the G-gene whereas figure-3A/B showed how the repetitive amino acids could shape the G-protein structure. Sadly, the 3-D structure of HMPV G-protein was not performed yet to know the role of different repetitive domains in 256aaG and 279aaG of HMPV. The SWISS-Model predicted alpha-helical structure of NH_2_-terminus few dozen amino acids due to its minimum 15% similarity with P2Y purinoreceptor 12 soluble cytochrome b562 (PDB ID: 4pxz). However, such study failed to predict any classification among 236aaG, 219aaG, 279aaG and 256aaG (data not shown)

In figure-4A, we demonstrated the multi-alignment of 229aa, 256aa and 279aa G-proteins to show the duplication or the deletion that occurred in A-genotypes. In figure-4B we have given a closest view of phylogeny with mixer of A, A1 and A2 or unknown as Ax. We consider here a dramatic change in nomenclature where A-genotype means 219aaG, A1-genotype means 279aaG with 180nt duplication while A2-genotype is 256aaG where is a 69nt deletion in the 279aaG HMPV genome. Surely, all scientists suggested the duplication with 180nt and 111nt in 279aaG and 256aaG proteins but our multi-alignment predicted the deletion theory.

Wei et al suggested that HMPV A-genotype peaked between 2000 and 2003, and then plateaued from 2003 to 2009. The A2b1 (G-180nt-dup) lineage most likely originated around 1992, whereas the A2b2 (G-111nt-dup) lineage probably originated from 1995. The A2b1-111-nt-dup sequences were derived from A2b1-180-nt-dup sequences as also we had found in our study. The evolutionary rate of HMPV genotype-A G gene was approximately 3.654× 10–3 substitution/site/year (95% highest probability density of 3.1303–4.0357) (Wei et al. 2023). The G-gene based classification was demonstrated in Spain where A2c genotype with 111nt duplication appeared to be more abundance during February-April whereas HMPV genome mutation rate suggested as 6.95x10^-4^ substitution/site/year (Pinana et al. 2023; Xie et al. 2021). Given to the high mutation rate individual amino acid changes in the G-protein between two genotypes or even in a same genotype sub-class, hard to pinpoint (figure-2B). However, our method had demonstrated the mechanism of G-protein diversity that was not found in other DNA/RNA viruses. The termination codon mutations in the G-gene were verified as demonstrated in figure-6A to figure-14A. The amino acid differences were clearly demonstrated in figure-6B to figure-14B.

The 5’-UTR of hMPV is short (5’-acgcgaaaaa aacgcgtata aattaagtta caaaaaaaca tgggacaagt gaaa-3’) while 3’-UTR is large 5’-gaaatgatga agatgataaa atagatgaca aattcatacc attctaaagt aattgtttga ttatgcaatt atatggtagt taattaaaa attaaaatta aaaatcaaa agttaaagtt taaaacctat cattaagttt attaaaaata agaaactat aattgaatgt atacggtttt tttgccgt-3’. The G>A and T>C mutations (red) were found with US clone at position 37 and 69 (PV052169) while another G>A mutation (green) was found at position 55 for Netherland A2-genotype clone (OL794464) and further one A>C mutation at position 32 (green) was detected in another Netherland A2-genotype clone (OL794360). As the 3’ and 5’ ends are incomplete in many clones, we have not studied the genotype variation in these regions but identical with China clone (OP358285), US clone (PP947648) and Japan clone (LC769215) (Nelson et al. 2023).

For diagnostic A/B genotyping, we found N-, F-and L-genes could be used after PCR sequencing of the genes. For N gene we found two oligos for A and B genotypes identification by BLASTN search. Npf1B-5’-gaagaaatagacaaagaggcaag-3’ 23nt oligo for A-genotype and Npf1A-5’-gaagagatagacaaagaagcaag-3’ 23nt oligo for B-genotype could be easily used. Similarly, for L-gene we found two 33nt oligos: Lpf1A-5’-CCT ATA GAC ATG CAC CAC CAG AAA CAA AAG GTG-3’ for A-genotype and Lpf1B-5’-CAT ACA GAC ATG CAC CAC CAG AAA CAA AAG GGG -3’ for B-genotype.

Scientists are using conserved F-gene genotyping today. Galiano M et al first showed that F protein was highly conserved in Argentinean patients, whereas the G protein showed extensive diversity between groups A and B (Galiano et al. 2006). Similar study by Arnott A et al showed 56.3% and 34.2% homology at the nucleotide and amino acid levels of G protein, respectively. A Japan study using F-gene PCR and multi-alignment suggested A2-genotype prevalence and no substitution in the gene as compared F-protein homology with other countries (Nidaira M et al. 2012). The F gene requires for fusion and attachment of the virus (Cox et al. 2013). Hindupur et al performed phylogenetic analysis of the partial F-gene of HMPV and revealed the presence of A2c subcluster among the study Indian strains as well as with B1 and B2 lineages (Hindupur et al. 2022). Our study will reinforce scientists to stick on G-gene based genotyping. Such study will help to emerge the basis of immunogenicity differences and attachment to host cells among the 219aaG, 236aaG, 256aaG and 279aaG proteins. The US data clearly showed the emergence of 256aaG lineages in the US patients in 2020 (Washington University study). The recombination gene was not pinpointed in HMPV. Thus, host recombination protein may be involved. The RdRP gene has mere similarity to COVID-19. But such minor similarity to spike protein of SARS-CoV-2 and HMPV RdRP was found (data not shown). We found an insignificant homology of F-protein with small accessory proteins (nsp7, nsp8, nsp10 and nsp9) of SARS-CoV-2 and proposed a mechanism of proteolytic processing of F-protein in vivo. The human RSV virus biology remains the main pathways of hMPV virus and more study needed for drug design and vaccine development (Yu et al. 2021; Meanwell et al. 2005).

## Conclusion

We have given a clear understanding of G-gene duplication and deletion as well as termination codon mutation in A1/A2 and B1/B2 genotypes creating varying lengths of G-proteins with varying amino acid sequence in the carboxy terminus. Thus, we revamped the G-gene genotyping to address A1/A2 and B1/B2 genotypes as well as the amino acid mutations among sub-genotypes A2a, A2b and A2c. We put important WGS information of HMPV in three tables so that fee software-based genetic analysis (BLASTN, BLASTP and CLUSTAL-Omega multi-alignment) of hMPV become easy and correct. Finally, we have given a new method of G-gene genotyping. The G-protein heterogeneity and antigenic variation is an important aspect of new population of HMPVs. Surely, our method of genotyping of HMPV sequences will encourage scientists to mention the genotype during their GenBank submission. The AMPV and HRSV biology must be studied with HMPV to tract the genetic changes during evolution in correlation with pneumonia severity among children to reduce the child mortality.

## Acknowledgement

The author thanks NCBI GenBank database and BLAST search engine for free service worldwide. The free MultAlin Software, CLUSTAL-Omega software and SWISS-Model program also acknowledged. AKC is retired professor of Biochemistry.

## Competent Interest

The author declares no competing interest

## Ethical Issues

The data presented here was computer generated and no animal or human was used.

